# Proteomic profiling of UV damage repair patches uncovers histone chaperones with central functions in chromatin repair

**DOI:** 10.1101/2024.08.23.609352

**Authors:** Alexandre Plessier, Audrey Chansard, Eliane Petit, Julia Novion Ducassou, Yohann Couté, Sophie E. Polo

## Abstract

DNA damage compromises not only genome stability but also the integrity of the chromatin template, which plays a central role in controlling cell identity. Our understanding of chromatin repair mechanisms is very incomplete. To bridge this knowledge gap, here we devise a novel proteomic strategy to characterize dynamic changes in the chromatin landscape during the repair of UV-induced DNA lesions in human cells, in a quantitative, unbiased and time-resolved manner. Thus, we identify the histone chaperones DNAJC9 and MCM2 as central players in chromatin repair. We demonstrate that DNAJC9 and MCM2 are independently recruited to sites of UV damage repair. DNAJC9 provides new H3-H4 histones to CAF-1 and HIRA chaperones for deposition into chromatin and also stimulates old H3-H4 histone recovery. DNAJC9 cooperates with MCM2 to coordinate old and new histone dynamics during UV damage repair. Together, our proteomic dataset provides a molecular framework for further dissecting epigenome maintenance mechanisms.

## INTRODUCTION

Chromatin is made of DNA wrapped around histone proteins in the cell nucleus and provides epigenetic information that governs cell identity by modulating gene expression (1). This pivotal information encompasses multiple layers of regulation, including DNA and histone modifications (2, 3) as well as histone variants (4), which confer specific structural and functional features to the so-called epigenome. Chromatin is in a dynamic state owing to histone deposition and eviction by histone chaperones (5), which can be assisted by chromatin remodelers (6) in promoting histone dynamics. Histone post-translational modifications further contribute to chromatin plasticity via dedicated factors that deposit, erase or read these marks (2, 7).

Epigenome stability is critical for the maintenance of cell function and homeostasis, as underlined by epigenome alterations in disease (8, 9). Nevertheless, maintaining epigenome stability is a challenge as all DNA metabolic activities entail some level of chromatin reorganization (10, 11). This challenge is particularly prominent during DNA damage repair because DNA damage can arise in an unscheduled manner at any time anywhere in the genome (12), which poses a major and unpredictable threat to both genome and epigenome stability. As such, chromatin plasticity is an integral part of the DNA damage response, DNA and chromatin repair being tightly coordinated (13).

Histone dynamics during DNA repair have been extensively characterized in response to UV damage in mammalian cells. Chromatin is transiently disrupted at UV damage sites with the eviction of H1 and H2A.Z histones (14, 15) while H3-H4 histones re-distribute away from damaged sites without dissociating from chromatin (16). This local reorganization of chromatin is thought to increase chromatin accessibility, thus facilitating the recruitment of DNA repair factors. DNA repair is accompanied by a restoration of chromatin through the recovery of parental H3-H4 histones (16) and the deposition of newly synthesized histones, as shown for H2A, H2A.X, H3.1 and H3.3 histone variants, all deposited de novo by specific histone chaperone complexes that act sequentially during the repair process. Deposition of new H3.3 histones by the histone chaperone complex HIRA occurs at early stages of UV damage repair, coupled to UV damage detection (17), while the de novo deposition of H3.1 and H2A variants by the CAF-1 and FACT chaperone complexes, respectively, take place during late repair steps, coupled to repair synthesis (15, 18, 19). Repaired chromatin is thus a mixture of original and new information, brought about by parental histones that characterize the pre-damage chromatin, and by newly synthesized histones, respectively.

Throughout the repair process, histone chaperones play a central role in controlling histone dynamics. These factors were identified through candidate approaches based on their cognate histone variants, which were shown to accumulate at sites of UV damage repair. We have now reached the limit of those approaches, and yet several gaps remain in our knowledge of chromatin dynamics during DNA repair. In particular, (i) it is unknown which factors provide new histones to the chaperones identified so far for de novo deposition at repair sites, (ii) the chaperones regulating parental histone redistribution and recovery are uncharacterized, and (iii) how the coordination of parental and new histone dynamics is achieved is still an open question. To fill these gaps, we have decided to adopt an unbiased strategy in order to identify new players in chromatin repair in an agnostic manner. We have devised a novel method for the analysis of proteins enriched at sites of UV damage repair in human cells with increased sensitivity and temporal resolution throughout the repair process. This method is based on the isolation of proteins on nascent DNA at repair sites (IPOND-R), combined with their characterization by mass spectrometry (MS)-based proteomics. With this method, we have identified histone chaperones that coordinate new H3-H4 deposition and parental H3-H4 recovery during chromatin repair.

## RESULTS

### The IPOND-R method captures proteins at sites of DNA damage repair

To profile new chromatin interactors during the repair of DNA damage in human cells, we devised a novel approach based on the established IPOND methodology (Isolation of Proteins On Nascent DNA) (20, 21), allowing us to specifically identify proteins associated with DNA repair patches in an unbiased, quantitative and time-resolved manner. We called this method IPOND-R for Isolation of Proteins On Nascent DNA at Repair sites (Figure 1A). In order to capture newly-synthesized DNA associated with DNA repair and not with DNA replication, IPOND-R includes a short-term treatment of cells with replication inhibitors, Hydroxyurea (HU) and Cytosine Arabinoside (Ara-C), for 2 h prior to DNA damage induction by global UVC irradiation. Cells are exposed to 10 J/m^2^ UVC, which generates UV lesions every 6 kb on average (22) and broadly distributed through the genome. Cells are then pulsed with Ethynyl-deoxyUridine (EdU) to label repair patches, crosslinked at different time points after UV irradiation and EdU is biotinylated through a Click-it reaction. After sonication of chromatin, proteins associated with biotinylated nascent DNA are pulled-down with streptavidin-coated beads for subsequent proteomic analyses (Figure 1A). Since there are no major variations in UV damage repair during the cell cycle, this method can be run on asynchronous cell populations.

**Figure 1:**
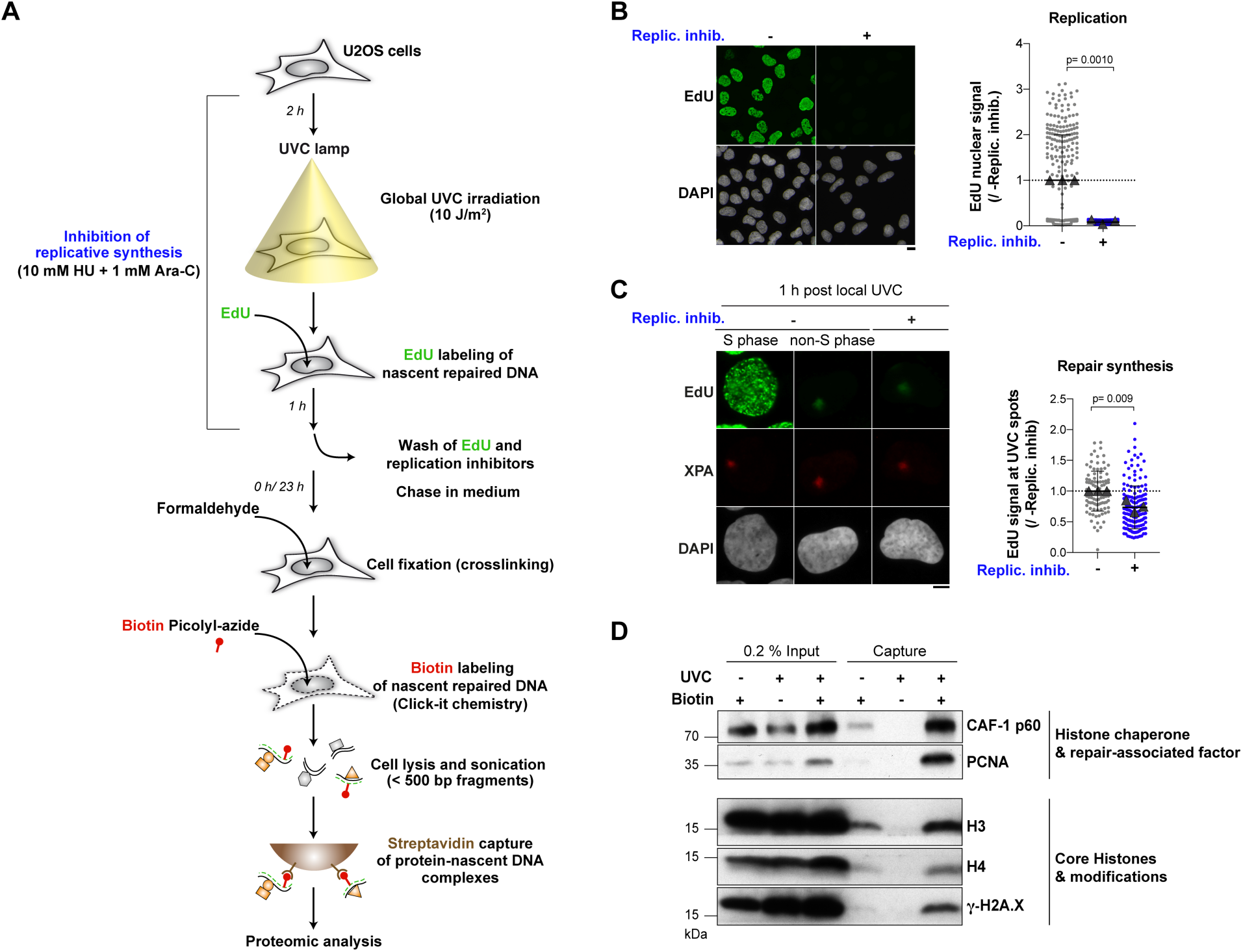
IPOND-R, a proteomics method for isolating proteins on nascent DNA at repair sites. (**A**) Scheme of the IPOND-R method. (**B**) Inhibition of replication-coupled DNA synthesis measured by EdU incorporation (pulse of 10 min) in U2OS cells treated for 2 h with the replication inhibitor cocktail (Replic. inhib.). (**C**) Effect of cell pre-treatment with the replication inhibitor cocktail (Replic. inhib.) on repair-coupled DNA synthesis measured by EdU incorporation (pulse of 1 h) at sites of local UV irradiation (marked by XPA immunostaining) in U2OS cells. S phase cells present in the non-treated population were identified by pan-nuclear EdU staining and excluded from the analysis. (**D**) Western blot analysis of proteins captured by IPOND-R 1 h post UV irradiation in U2OS cells. -UV and –Biotin are used as negative controls. Dot plots: mean ± SD from at least 99 cells scored in 3 independent experiments (the mean of each experiment is shown as a grey triangle). Statistical significance is calculated with a two-sided Student’s t-test with Welch’s correction. Scale bars, 10 μm.

We verified that short term cell treatment with the replication inhibitor cocktail was sufficient to erase replicative DNA synthesis (Figure 1B) while still allowing DNA repair synthesis at sites of UV damage, although with a decrease in efficacy (Figure 1C). In line with this, we could detect a transient inhibition of gap filling at UV damage sites upon replication inhibition as shown by increased RPA levels 1 h after local UV irradiation returning to normal 2 h post UV (Figure S1A). Cells treated with replication inhibitors also showed signs of replicative stress as illustrated by a transient increase in γH2A.X and phospho-Chk1 signals and an altered cell cycle distribution with an accumulation of cells in G1 24 h post UV (Figure S1B-D). Nevertheless, replication inhibition did not impair the recruitment of UV damage repair factors to damage sites (Figure S1E), and the kinetics of UV lesion removal (Figure S1F-G). Furthermore, we verified that cell treatment with replication inhibitors allowed efficient and timely recruitment of the histone chaperones CAF-1 and HIRA and subsequent deposition of newly synthesized H3.1 and H3.3 histone variants at UV damage sites (Figure S1H-K).

For EdU biotinylation, we used an enhanced version of biotin-azide conjugate called biotin picolyl azide, which improved in vivo biotinylation efficiency by 4.5-fold as verified by imaging with fluorescently-labeled streptavidin (Figure S2A). Note that DNA breakage arises during the biotinylation reaction due to copper-catalyzed hydrolysis of the DNA (21) but as this occurs after cell fixation, it does not introduce any significant bias in the analysis. UV irradiation led to a detectable increase in biotinylated EdU (Figure S2B), thus supporting the sensitivity of the proposed methodology to monitor repair synthesis. Importantly, even though the UV damage repair machinery can excise EdU (23), the UV-associated EdU signal was stable for up to 24 h post UVC irradiation in our experimental setting (Figure S2B), allowing us to perform long-term analyses. We also verified that a 1-h EdU pulse immediately after UV irradiation allowed the capture of repair events in heterochromatin regions, known to repair UV lesions more slowly than euchromatin (24, 25). For this, we employed a validated murine cell line model, NIH/3T3 cells stably expressing the DNA damage sensor DNA damage binding protein 2 (DDB2) (14), for studying UV damage repair in DAPI-dense pericentric heterochromatin domains. We observed that the relative levels of EdU in heterochromatin vs. euchromatin were comparable when we labeled early repair events (EdU pulse immediately after UV) or later repair events (EdU pulse 3 h after UV) (Figure S2C), indicating that the IPOND-R analysis is not biased towards repair in euchromatin regions.

As a proof-of-principle, we performed IPOND-R in human U2OS cells and analyzed by western-blot proteins associated with the repair patch 1 h post UV irradiation (Figure 1D). Core histones, the damage-induced histone modification γH2A.X, PCNA and the histone chaperone CAF-1 were pulled down only in the presence of biotin and enriched in the capture samples from UV-treated cells compared to non-irradiated cells, demonstrating that IPOND-R efficiently and specifically captures chromatin repair patches. Altogether, these results indicate that the IPOND-R method is suited for studying chromatin dynamics specifically at sites of DNA damage repair, in different chromatin contexts and throughout the repair process, from early to late repair steps.

### Identification of chromatin regulators enriched at sites of UV damage repair

In order to identify new players in chromatin repair, we combined IPOND-R with unbiased protein identification by mass spectrometry. We detected 255 proteins that were significantly enriched at repair patches 1 h post UV irradiation in U2OS cells with a +/-UV fold change > 2 and a limma p-value < 0.01 (Figure 2A, Table S1). Among these proteins, we identified known UV damage repair factors (XPC, DDB1, CUL4A, PCNA) as expected, supporting the validity of the IPOND-R method in capturing UV damage repair patches. There was a modest overlap with the dataset obtained by Stefos et al. (26) in human skin fibroblasts 4 h post UV (45 proteins in common) and we identified 197 proteins that were not found in previous proteomic studies post UV (Figure S3A) (26, 27). Much fewer proteins, 12, were retrieved as enriched at repair sites 24 h post UV irradiation and those were mostly core chromatin components (histones and high-mobility group proteins) (Figure 2B, Table S1). Comparing the 1 h and 24 h time points allows to eliminate factors constitutively bound to chromatin in order to focus on those that transiently bind to chromatin during repair (247 1 h-specific hits) (Figure 2C, Table S1). Gene ontology analysis of the 1 h-specific hits revealed an enrichment of biological processes associated with DNA and RNA metabolic activities, the nuclear periphery (nuclear pores, lamins), the immune system (isotype switching and somatic hypermutation), and chromatin organization (Figure 2D, S3B). To further study those chromatin regulators, we crossed our dataset with the EpiFactors database (Figure 2E). Histone modification activities (writers, readers, erasers) and histone chaperones were the most represented, followed by chromatin remodelers and DNA modifiers (Figure 2E-F). We validated the presence of the chromatin remodeler catalytic subunit SMARCA5 and of the histone modifying enzyme HDAC1 in the +UV capture by IPOND-R coupled to western blot detection (Figure 2G).

**Figure 2:**
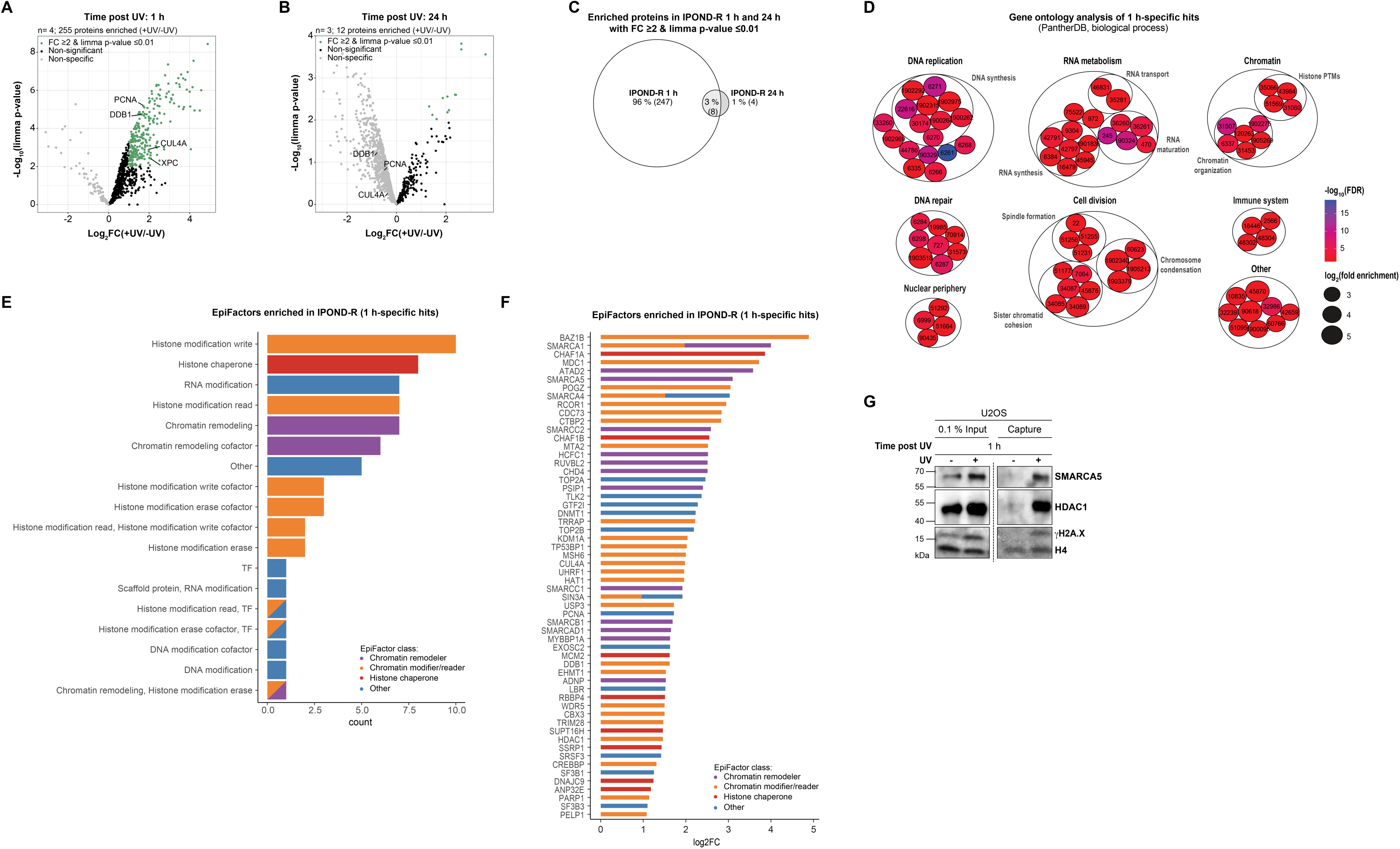
Identification of factors enriched at sites of UV damage repair by IPOND-R. (**A-B**) Volcano plots showing mass spectrometry results of IPOND-R experiments performed at 1 h (A) and 24 h (B) post global UVC in U2OS cells (n= number of independent experiments). Proteins significantly enriched in irradiated compared to non-irradiated samples (+/-UV fold-change (FC)>= 2 and limma p-value<= 0.01) are shown in green, non-significant proteins are shown in black, and proteins with a negative fold-change (non-specific) are shown in grey. Nucleotide excision repair proteins reaching significance at 1 h are highlighted. (**C**) Venn diagram showing enriched proteins at 1 h and 24 h post UV damage with a +/-UV fold-change (FC)>= 2 and limma p-value<= 0.01. (**D**) Gene ontology analysis of the 1 h-specific hits detected by IPOND-R. Biological process terms with a log2 enrichment>= 2,8 and - log10(False Discovery Rate)<= 2 are shown with their reference number and grouped based on their association with DNA replication, DNA repair, nuclear periphery, RNA metabolism, cell division, chromatin, immune system, or other. The - log10-False Discovery Rate is represented by the color, and the log2-fold enrichment of the biological processes by the circle size. (**E**) Bar plot showing the enrichment of EpiFactor classes in the 1h-specific IPOND-R dataset colored by categories: chromatin remodelers in purple, chromatin modifiers in orange, histone chaperones in red (TF, transcription factors). (**F**) Bar plot showing the EpiFactors in the 1 h-specific IPOND-R dataset ranked by +/-UV fold-change (FC) and colored by categories: chromatin remodelers in purple, chromatin modifiers in orange, histone chaperones in red. SMARCA5 and HDAC1, known EpiFactors recruited to UV-damaged chromatin, are highlighted and validated by Western Blot in (G). (**G**) Western blot validation of the EpiFactors HDAC1 and SMARCA5 captured by IPOND-R 1h post UV irradiation in U2OS cells. -UV is used as negative control.

Among histone chaperones, we retrieved most of those known to be involved in chromatin repair following UV damage (Figure 3A), with the three subunits of the CAF-1 complex (CHAF1A, CHAF1B, RBBP4), which deposits new H3.1 (18), both subunits of the FACT complex (SUPT16H, SSRP1), which promotes new H2A.X deposition and ANP32E, which promotes H2A.Z eviction in UV-damaged chromatin (15). In addition, we also detected additional histone chaperones with yet unknown functions in chromatin repair (Figure 3A), namely Minichromosome Maintenance Complex Component 2 (MCM2) and DnaJ Heat Shock Protein Family Member C9 (DNAJC9). MCM2 was shown to chaperone H3-H4 histones at replication forks (28, 29) and the heat-shock protein DNAJC9 (30) was recently characterized as a H3-H4 histone chaperone in human cells (31, 32). Notably, our optimized IPOND-R methodology permits the unprecedented detection of DNAJC9 (+/-UV fold change= 2.36; limma p-value= 1.53E-3 at 1 h post UV) and MCM2 (+/-UV fold change= 3.07; limma p-value= 6.16E-4 at 1 h post UV) at sites of UV damage repair while these chaperones were not identified in previous proteomic analyses post UV (26, 27). All the histone chaperones that we identified on UV damaged chromatin were captured at 1 h and not 24 h post UV (Figure 3A-B), highlighting their dynamic recruitment to chromatin during the repair process. We validated these findings by western blot on the capture material in U2OS cells (Figure 3C) and in non-cancerous RPE-1 cells (Figure 3D), which confirmed the transient recruitment of DNAJC9 to repair sites. We further validated DNAJC9 and MCM2 recruitment to sites of UV damage repair through an orthogonal approach that does not rely on using replication inhibitors. For this, we imaged these histone chaperones by immunofluorescence in cells exposed to local UVC irradiation. First, we assessed DNAJC9 enrichment at sites of UVC laser-induced damage in U2OS cells expressing a GFP-tagged form of the UV damage sensor protein DDB2. DNAJC9 showed a distinct and significant enrichment of about 2-fold at UV damage sites (Figure 3E). We confirmed this result by irradiating cells with a UVC lamp through micropore filters and observed a similar albeit more modest enrichment of DNAJC9 and MCM2 at UV damage sites (1.2-fold, Figure 3F-G). Collectively these findings highlight the potential of IPOND-R in identifying novel chromatin-associated factors at sites of UV damage repair, including the histone chaperones DNAJC9 and MCM2.

**Figure 3:**
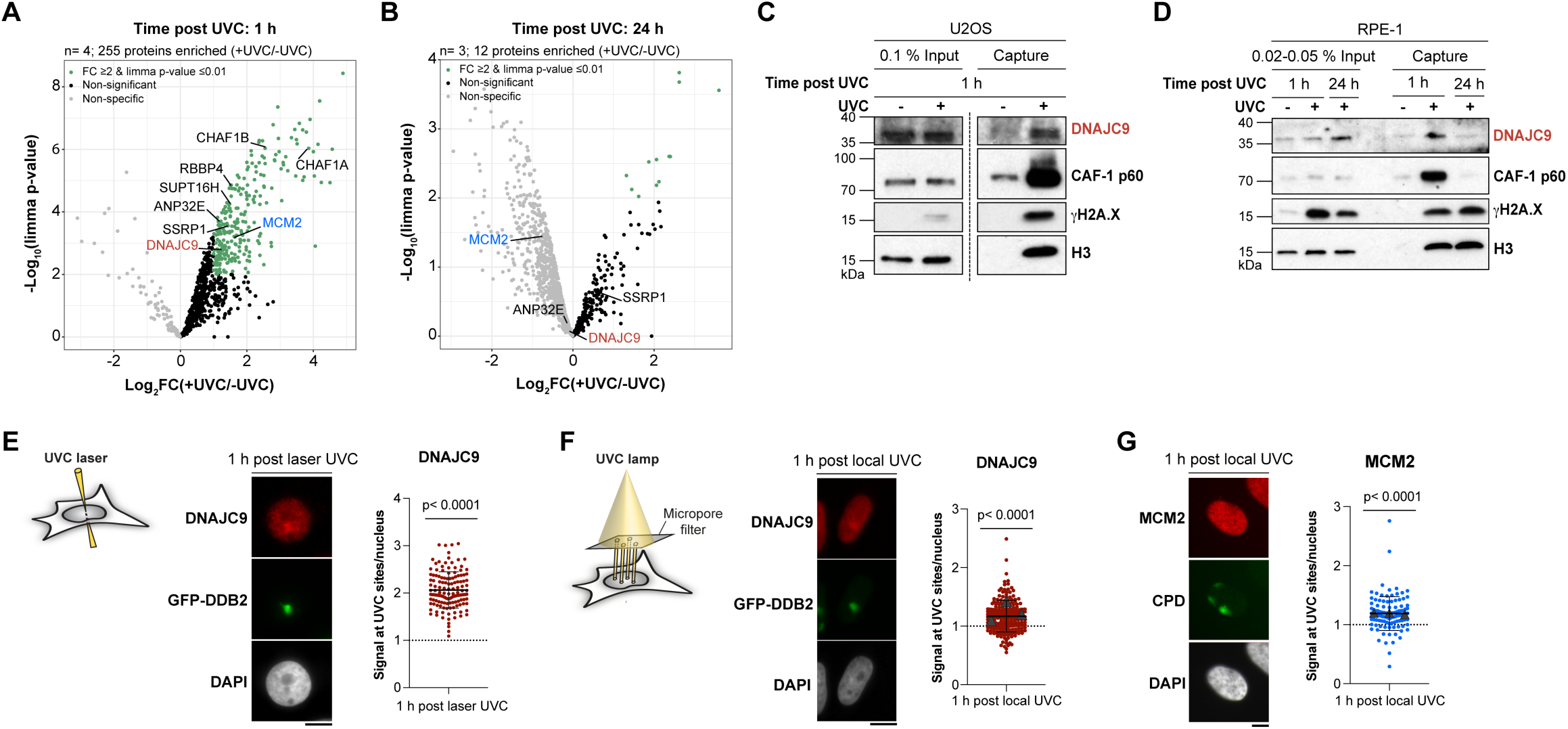
DNAJC9 and MCM2 histone chaperones are recruited to sites of UV damage repair. (**A-B**) Volcano plots showing mass spectrometry results of IPOND-R experiments performed at 1 h (A) and 24 h (B) post global UVC in U2OS cells as in Figure 2. Known histone chaperones reaching significance at 1 h are highlighted. (**C**) Western blot validation of some histone chaperones (DNAJC9 and the CAF-1 subunit p60) captured by IPOND-R 1h post UVC irradiation in U2OS cells. -UV is used as negative control. (**D**) Western blot showing the dynamic recruitment of DNAJC9 and CAF-1 p60 to sites of UV damage repair in RPE-1 cells by IPOND-R at 1 h and 24 h post UV damage. -UV is used as negative control. (**E**) Immunodetection of DNAJC9 1 h post UVC laser micro-irradiation in U2OS cells stably expressing GFP-DDB2 (UV damage sensor). The enrichment of DNAJC9 at UV damage sites relative to the nucleus is shown on the graph. (**F**-**G**) Immunodetection of DNAJC9 (F) and MCM2 (G)1 h post local UVC irradiation through micropore filters in U2OS cells. The enrichment of these chaperones at UV damage sites (marked by GFP-DDB2 or CPD) relative to the nucleus is shown on the graphs. Dot plots: mean ± SD from at least 110 cells scored in 1-3 independent experiments (the mean of each experiment is shown as a grey triangle). Statistical significance is calculated with a one sample t-test compared to a theoretical mean of 1. Scale bar, 10 μm.

### The histone chaperone DNAJC9 is not involved in DNA repair following UV damage

To study the function of DNAJC9 and MCM2 in the DNA damage response, we first focused on DNAJC9 as it was the least characterized histone chaperone. We employed siRNA-mediated depletion because DNAJC9 knock-out was reported to be lethal in human cells (31, 32), and we reached an effective depletion of DNAJC9 of over 50% in U2OS cells (Figure S4A). Since DNAJC9 was recruited early during the DNA damage repair process, we addressed whether this histone chaperone may participate, directly or indirectly, to the repair of DNA damage. However, we did not observe any significant effect of DNAJC9 depletion on the recruitment of the UV-damage repair factor XPB, involved in the unwinding of the DNA double-helix prior to damage excision (Figure S4B), nor on DNA repair synthesis, monitored by EdU incorporation at UV damage sites (Figure S4C). Furthermore, the kinetics of UV photoproduct removal (CPD and 6,4-PP) were also unchanged upon DNAJC9 knock-down while depletion of the repair factor XPG prevented UV lesion removal (Figure S4D). In line with this, DNAJC9 depletion did not impair the survival of cells to UV irradiation (Figure S4E). Taken together, these results rule out a function for DNAJC9 in the repair of DNA damage and prompted us to explore a direct role of this chaperone in histone dynamics at sites of UV damage repair.

### DNAJC9 promotes the deposition of newly synthesized H3 variants at sites of UV damage repair

Given the H3-H4 histone binding activity of DNAJC9, we investigated whether this histone chaperone may impact the deposition of newly synthesized H3 histone variants at UV damage sites. We employed the SNAP-tag technology to track new H3.1 and H3.3 histones in U2OS cells stably expressing SNAP-tagged H3 variants (33) and monitored their accumulation at sites of local UVC irradiation upon depletion of DNAJC9 (Figure 4A). Given that H3.1 is also de novo deposited at replication foci (34), the incorporation of new H3.1 histones was analyzed outside S-phase to focus on repair-coupled deposition. We observed a marked and reproducible decrease of new H3.1 and H3.3 levels at sites of UV damage upon DNAJC9 depletion (Figure 4B-C), which we confirmed with a second siRNA targeting DNAJC9 (Figure S5A-B). We also recapitulated these results in non-cancerous RPE-1 cells stably expressing SNAP-tagged H3 variants (Figure S5C-D). Contrary to what observed with H3 histone variants, DNAJC9 depletion did not affect the de novo deposition of the H2A.X variant at UV damage sites (Figure S5E), which is fully consistent with the H3-H4 specificity of this histone chaperone (31, 32). Importantly, DNAJC9 depletion did not affect the total protein levels of the histone chaperones CAF-1 and HIRA (Figure S5A-B), known to promote the de novo deposition of H3.1 and H3.3 in UV-damaged chromatin (Figure 4D), and also did not impair the recruitment of these chaperones to UV damage sites (Figure 4E-F). From these results, we conclude that DNAJC9 promotes new H3 variant deposition at sites of UV damage repair without affecting their cognate histone chaperones CAF-1 and HIRA.

**Figure 4:**
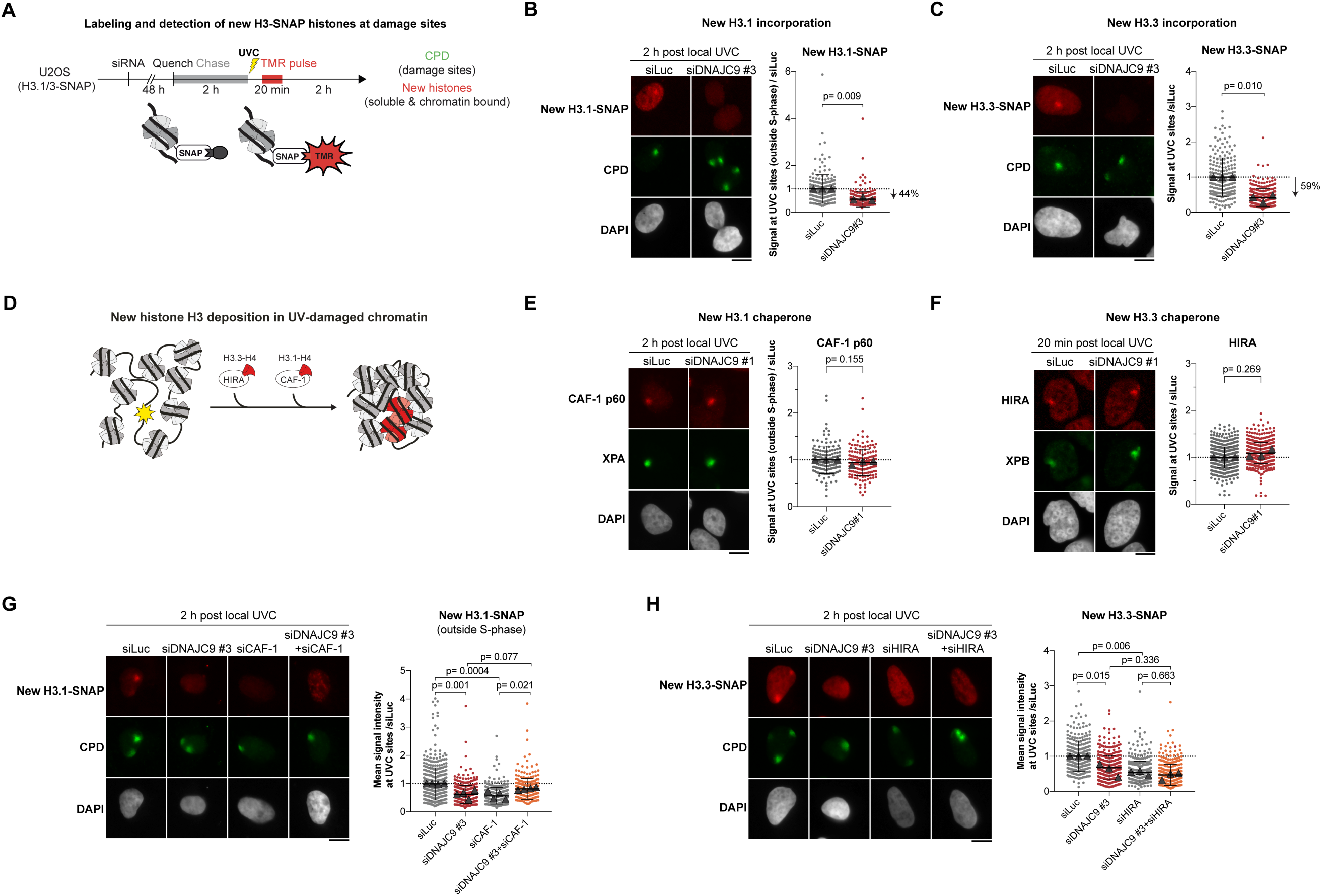
DNAJC9 stimulates new H3 deposition at sites of UV damage repair. (**A**) Protocol for new H3-SNAP histone detection at UV damage sites. (**B-C**) Detection and quantification of newly synthesized H3.1-SNAP (B) and H3.3-SNAP (C) histones at sites of UVC irradiation (marked by CPD) in U2OS cells stably expressing SNAP-tagged H3 variants and treated with the indicated siRNAs (siLuc, control). The incorporation of new H3.1-SNAP histones is quantified in cells outside S-phase, S-phase cells showing a typical focal pattern of new H3.1 deposition throughout the nucleus. The percentages of signal decrease observed in siDNAJC9-treated cells are indicated on the right. (**D**) Histone chaperones (white) promoting the deposition of newly synthesized H3 variants (red) in UV-damaged chromatin (yellow star, UV lesion). (**E-F**) Immunodetection of CAF-1 (E) and HIRA (F) chaperones at sites of UVC irradiation (marked by XPA and XPB, respectively) in U2OS cells treated with the indicated siRNAs (siLuc, control). CAF-1 signal at damage sites is quantified in non-S phase cells. (**G-H**) Detection and quantification of newly synthesized H3.1-SNAP (G) and H3.3-SNAP (H) histones at sites of UVC irradiation (marked by CPD) in U2OS cells stably expressing SNAP-tagged H3 variants and treated with the indicated siRNAs (siLuc, control). The incorporation of new H3.1-SNAP histones is quantified in cells outside S-phase. Dot plots: mean ± SD from at least 155 cells scored in 3-5 independent experiments (the mean of each experiment is shown as a grey triangle). Statistical significance is calculated with a two-sided Student’s t-test with Welch’s correction (B-C, E-F) or one-way ANOVA with multiple comparisons (G-H). Scale bars, 10 μm.

In addition to a role at damage sites, we suspected that DNAJC9 may also promote new histone deposition in undamaged chromatin, during replication in particular, because this histone chaperone sustains cell cycle progression into S phase (Figure S5F), which is consistent with its reported function in cell proliferation (31). Similar to what was observed at UV damage sites, DNAJC9 depletion impaired the incorporation of newly synthesized H3.1 histones at replication foci in S phase and also the replication-independent deposition of H3.3 in undamaged conditions (Figure S5G-H). These results highlight a functional role of DNAJC9 in chromatin assembly in vivo via the deposition of new H3.1 and H3.3 histones, both in basal conditions and in response to DNA damage.

We next sought to characterize whether DNAJC9 may function in the same histone deposition pathways as the CAF-1 and HIRA chaperones at repair sites. To do so, we compared single and double knock-downs of DNAJC9 and CAF-1 or HIRA (Figure S5I-L) for their impact on new H3 histone deposition at UV damage sites (Figure 4G-H). Single depletions of DNAJC9 and CAF-1 similarly impaired the incorporation of new H3.1 histones at damage sites (Figure 4G) and single depletions of DNAJC9 and HIRA had comparable effects on new H3.3 deposition at UV sites (Figure 4H). The impact on new H3 variant deposition was partial in each condition, and we did not observe any additive effect upon double depletion of DNAJC9 and the other chaperones (Figure 4G-H). These results establish that DNAJC9 functions in the same histone deposition pathways as CAF-1 and HIRA to promote the incorporation of new H3.1 and H3.3 histones at sites of UV damage repair.

### DNAJC9 promotes the recovery of parental H3 histones at sites of UV damage repair

We next investigated whether, besides its activity in promoting new histone deposition at repair sites, DNAJC9 may also control the recovery of old H3-H4 histones that redistribute away from damaged chromatin regions (16). The only factor known so far to control parental histone dynamics at UV sites is the UV damage sensor protein DDB2, whose binding to damaged chromatin and subsequent release promote the redistribution and recovery of parental histones, respectively (16). To test DNAJC9 contribution to these dynamics, we used the SNAP-tag technology to mark old H3.3 histones in U2OS cells stably expressing H3.3-SNAP, and quantified their levels at UV damage sites compared to the entire nucleus (Figure 5A). The depletion of old H3.3 histones at UV damage sites early after irradiation (10 min) and their subsequent enrichment at late time points (8 h) reflect parental histone redistribution and recovery, respectively (Figure 5B, siLuc condition). This recovery was impaired when DNAJC9 was depleted, similar to what was observed upon depletion of the late repair factor XPG, used as a positive control (Figure 5B). These results demonstrate that DNAJC9 promotes parental H3.3 histone recovery at sites of UV damage repair. To test whether this may result from impaired DDB2 release from chromatin, we examined DDB2 dynamic binding to UV-damaged chromatin in the absence of DNAJC9 in U2OS cells stably expressing GFP-tagged DDB2. While XPG depletion prevented DDB2 release by inhibiting DNA repair progression, as expected, the depletion of DNAJC9 had no effect on DDB2 release from chromatin (Figure 5C). Together, these findings demonstrate that the histone chaperone DNAJC9 promotes parental H3 histone recovery at repair sites independently of the UV damage sensor DDB2. Consistent with a role of DNAJC9 in old histone recycling 8 h post UV irradiation, we detected the presence of this histone chaperone at sites of UV damage 8 h post irradiation by immunofluorescence (Figure 5D). Given that DNAJC9 promotes both new histone deposition and old histone recycling at UV damaged sites, we reasoned that DNAJC9 may control the overall histone density at repair sites. We used the SNAP-tag technology to mark all H3.3-SNAP histones 8 h post irradiation (Figure 5E) and observed a modest but significant decrease in the total levels of H3.3-SNAP histones at UV sites when DNAJC9 was depleted (Figure 5F-G). Therefore, DNAJC9 maintains histone density at sites of UV damage repair through new histone deposition and old histone recycling.

**Figure 5:**
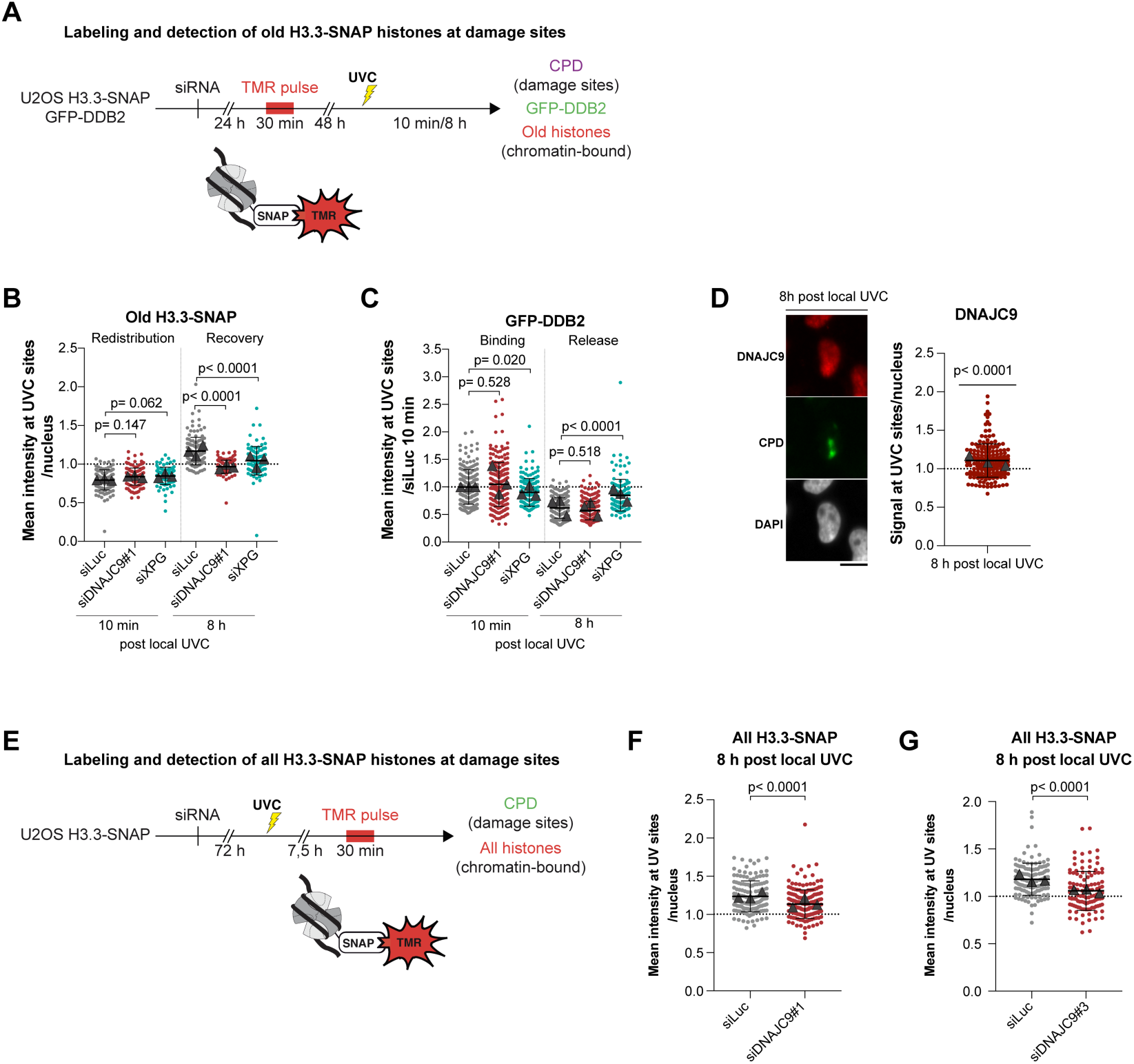
DNAJC9 promotes old histone H3 recovery during chromatin repair. (**A**) Protocol for old H3.3-SNAP histone detection at UV damage sites. (**B**) Detection and quantification of old H3.3-SNAP histone enrichment at sites of UVC irradiation (marked by CPD) relative to the nucleus in U2OS cells stably expressing SNAP-tagged H3.3 and treated with the indicated siRNAs (siLuc, negative control; siXPG, positive control inhibiting old histone recovery). (**C**) GFP-DDB2 levels at UVC damage sites (marked by CPD) measured in U2OS GFP-DDB2 cells treated with the indicated siRNAs (siLuc, negative control; siXPG, positive control inhibiting DDB2 release). (**D**) Immunodetection of DNAJC9 8 h post local UVC irradiation with a UVC lamp through micropore filters in U2OS cells. The enrichment of DNAJC9 at UV damage sites (marked by CPD) relative to the nucleus is shown on the graph. (**E**) Protocol for total H3.3-SNAP histone detection at UV damage sites. (**F-G**) Detection and quantification of total H3.3-SNAP histones at sites of UVC irradiation (marked by CPD) in U2OS cells stably expressing SNAP-tagged H3.3 and treated with siDNAJC9#1 (F) and siDNAJC9#3 (G) (siLuc, negative control). Dot plots: mean ± SD from at least 99 cells scored in 3 independent experiments. Statistical significance is calculated by one-way ANOVA with multiple comparisons (B-C), a two-sided Student’s t-test with Welch’s correction (F) or a one sample t-test compared to a theoretical mean of 1 (D). Scale bars, 10 μm.

### DNAJC9 cooperates with MCM2 to coordinate old and new histone dynamics at sites of UV damage repair

We next investigated how DNAJC9 may promote old histone recovery at repair sites. We envisioned that DNAJC9 could cooperate with the MCM2 histone chaperone in this process because MCM2 is engaged in histone-dependent interactions with DNAJC9 (32), and promotes the recycling of parental H3-H4 histones at replication forks (28). We first tested by immunofluorescence in cells exposed to local UVC irradiation whether MCM2 was still enriched on damaged chromatin at the time of old histone recovery. MCM2 enrichment at UV damage sites was detectable 8 h post irradiation and even more pronounced than at 1 h post irradiation (Figure 6A, siLuc conditions). MCM2 recruitment to sites of UV damage repair was unaffected by DNAJC9 knock-down and MCM2 depletion did not impair DNAJC9 enrichment at repair sites (Figures 6A-B), arguing that both histone chaperones are independently recruited to UV damage sites. We next tested if MCM2 could function together with DNAJC9 in repair-coupled histone dynamics. Similar to DNAJC9 depletion, MCM2 depletion by siRNA impaired old H3.3 recovery at sites of UV damage repair (Figure 6C) and the dual knock-down of both chaperones did not show any additive effect (Figures 6D and S6A). Thus, MCM2 cooperates with DNAJC9 to promote old histone recovery at repair sites. We obtained comparable results when examining new H3.3 deposition at earlier times following UV irradiation, arguing that MCM2 also cooperates with DNAJC9 to stimulate new histone deposition at sites of UV damage repair (Figures 6E and S6B). From these results, we conclude that the DNAJC9 and MCM2 histone chaperones play a central function in chromatin repair by coordinating old and new histone dynamics.

**Figure 6:**
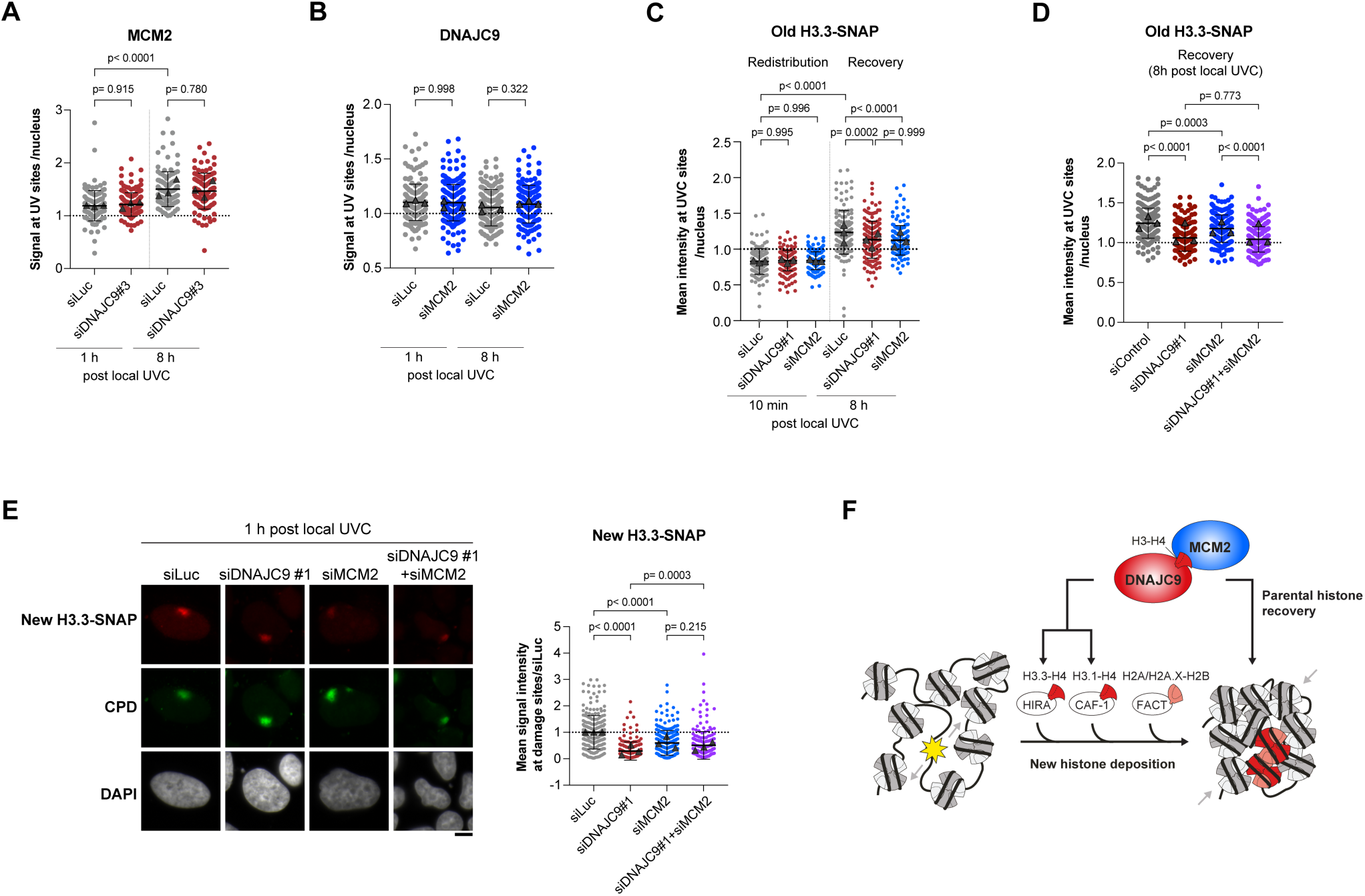
DNAJC9 and MCM2 cooperate to stimulate new histone deposition and parental histone recovery at DNA repair sites. (**A**) Immunodetection of MCM2 1 h and 8 h post local UVC irradiation in U2OS cells treated with the indicated siRNAs (siLuc, control). The enrichment of MCM2 at UV damage sites (marked by CPD) relative to the nucleus is shown on the graph. The first condition (siLuc, 1 h post UVC) is also presented in Figure 3G. (**B)** Immunodetection of DNAJC9 1 h and 8 h post local UVC irradiation in U2OS cells treated with the indicated siRNAs (siLuc, control). The enrichment of DNAJC9 at UV damage sites (marked by CPD) relative to the nucleus is shown on the graph. (**C-D**) Detection and quantification of old H3.3-SNAP histone enrichment at sites of UVC irradiation (marked by CPD) relative to the nucleus in U2OS cells stably expressing SNAP-tagged H3.3 and treated with the indicated siRNAs (siLuc, negative control) at 10 minutes and 8 h post irradiation (C) or at 8 h only (D). (**E**) Detection and quantification of newly synthesized H3.3-SNAP histones at sites of UVC irradiation (marked by CPD) in U2OS cells stably expressing SNAP-tagged H3.3 and treated with the indicated siRNAs (siLuc, control). (**F**) Working model. The histone chaperones DNAJC9 and MCM2 are recruited to sites of UV damage repair and jointly coordinate old and new histone H3-H4 dynamics during chromatin repair. Dot plots: mean ± SD from at least 104 cells scored in 3 independent experiments. Statistical significance is calculated by one-way ANOVA with multiple comparisons. Scale bars, 10 μm.

## DISCUSSION

Preserving chromatin integrity together with genome stability in response to genotoxic stress is key for safeguarding cell identity and protecting cells from pathological development. Yet, our understanding of chromatin repair mechanisms is still very incomplete. Here, we bridge this knowledge gap and identify new players in these mechanisms through the proteomic profiling of DNA repair patches.

While similar methods for labeling sites of DNA damage repair in non-replicating cells have been employed in recent works (26, 35), our proteomic approach presents several notable improvements compared to previous proteomic studies in UV-damaged cells (26, 27): (i) the present approach is focused on repair sites rather than a global analysis of the chromatin-bound proteome post damage infliction, (ii) our method is time-resolved with access to the whole repair process, from early to late repair steps, allowing us to discriminate proteins that dynamically bind to chromatin in response to damage from constitutively bound proteins, and (iii) the increased sensitivity of our methodology allows us to detect low abundant or weakly bound proteins that were not identified in previous datasets.

Among them, we focused our attention on the recently characterized histone chaperone DNAJC9 (31, 32). Thus, we uncover a function for DNAJC9 in chromatin assembly in vivo, which was previously characterized only in vitro (32). We hypothesize that DNAJC9 provides new H3-H4 histones to CAF-1 and HIRA chaperones for deposition into chromatin because DNAJC9 is loosely bound to chromatin, as it is mostly solubilized by detergents (data not shown), in contrast to CAF-1 and HIRA, suggesting an upstream function of DNAJC9 in the histone supply chain. Whether DNAJC9 directly provides histones to CAF-1 and HIRA is still unclear as the latter chaperones were not systematically identified as DNAJC9 interactors in basal conditions (32, 36, 37). By ensuring H3-H4 histone supply, DNAJC9 prevents mislocalization of the centromeric H3 variant CENP-A to chromosome arms (38). It is thus tempting to speculate that DNAJC9 may similarly prevent ectopic CENP-A deposition at sites of UV damage repair.

In addition to the reported function of DNAJC9 in controlling histone protein folding in the cytoplasm (31, 32), our work highlights a nuclear function of DNAJC9 in coordinating old and new H3-H4 histone dynamics during the repair process. Indeed, we show that DNAJC9 cooperates with MCM2 to stimulate both new histone deposition and old histone recovery during repair. Our discoveries not only reveal a function of these histone chaperones in chromatin repair but also instruct us on their activity and central position in the histone supply chain, at the crossroads of new histone deposition and old histone recycling (Figure 6F). The mechanism whereby DNAJC9 and MCM2 stimulate parental histone recovery during UV damage repair warrants further investigation.

Several pieces of evidence also suggest that DNAJC9 may coordinate new and parental histone dynamics in contexts other than DNA repair. Indeed, DNAJC9 is found enriched on nascent replicated chromatin (39) and we show that DNAJC9 stimulates new H3.1 deposition at undamaged replication forks. A potential role of DNAJC9 in parental histone recycling at replication forks, possibly in coordination with MCM2 (28), deserves to be investigated.

How MCM2 and DNAJC9 are recruited to sites of UV damage repair is not yet elucidated. It is tempting to speculate that their recruitment may be mediated by direct or indirect interactions with UV damage repair factors or with other histone chaperones enriched at repair sites. In this respect, MCM2 is involved in histone-dependent interactions with the FACT histone chaperone (36) that is recruited to UV sites coupled to repair synthesis (15). An additional candidate that could promote DNAJC9 recruitment to UV-damaged chromatin is RAD23A, which was identified in the DNAJC9 interactome in human cells (31) and binds the early UV damage repair factor XPC (40, 41). UV-damage responsive post-translational modifications of these histone chaperones may also regulate their function in chromatin repair (27). Our proteomic approach does not distinguish between repair in euchromatin and heterochromatin. However, we envision that histone chaperones may play distinct roles in different chromatin compartments. The use of murine cellular models suited for the analysis of heterochromatin repair of UV lesions (14) will be instrumental to address this issue.

We do not rule out the possibility that DNAJC9 and MCM2 may respond to other types of DNA lesions than UV photoproducts, and thus play a more general role in maintaining epigenome stability following genotoxic stress. Interestingly, a fungal ortholog of DNAJC9, Dnaj4, which also exhibits H3-H4 histone chaperone activity, was shown to safeguard genome integrity at least in part through the control of gene expression in response to DNA damage (42). Together with our findings, this hints towards an evolutionarily conserved function of DNAJC9 in genome and epigenome maintenance. DNAJC9 is a dual heat shock molecular and histone chaperone that recruits heat shock proteins of the HSP70 family via its J domain to promote histone protein folding (31, 32). Whether DNAJC9 functions in chromatin repair depend on binding to HSP70 family proteins remains to be investigated, which could be done using separation of function mutants in the J domain and the histone binding domain of DNAJC9 (32). Notably, HSPA4/HSP74 is significantly enriched at sites of UV damage repair in the 1 h IPOND-R dataset and thus represents a potential candidate that could cooperate with DNAJC9 in controlling histone dynamics during repair.

While in this study we focused on the histone chaperones DNAJC9 and MCM2, our proteomic analysis uncovered a number of additional interesting candidates as potential new players in chromatin dynamics during DNA repair. Among them, several histone modifiers and readers highlight a fine-tuned regulation of histone modifications at repair sites, which deserves to be investigated in future studies. The histone modifiers and readers identified in our dataset and by others (26, 27) will provide clues to the underlying mechanisms. Along the same lines, the DNA modifiers enriched at sites of UV damage repair will help characterizing the machinery for repair-coupled DNA methylation maintenance, thus broadening our understanding of epigenome maintenance mechanisms in response to DNA damage. Intriguingly, several nuclear pore components and lamins were also enriched at sites of UV damage repair, as seen in a previous study (26). There is growing evidence for a role of the nuclear periphery in the repair of DNA double-strand breaks, with some DSBs relocalizing to the nuclear periphery for repair and nuclear pores and lamina-associated chromatin domains regulating DSB repair pathway choice (43–46). However, the connection of UV damage repair with the nuclear periphery is still elusive and will be interesting to explore to further dig into how nuclear architecture may govern genome maintenance and reciprocally. Overall, our proteomic dataset provides a molecular foundation for further exploration of the fundamental mechanisms underpinning chromatin plasticity and integrity following genotoxic stress.

## MATERIAL AND METHODS

### Cell culture

U2OS (ATCC HTB-96, human osteosarcoma, female), RPE-1 hTERT (ATCC CRL-4000, human retinal pigment epithelial cell, female), and NIH/3T3 cells (ATCC CRL-1658, mouse embryonic fibroblast, male) were grown at 37°C and 5% CO2 in Dulbecco’s modified Eagle’s medium (DMEM, Invitrogen) supplemented with 10% fetal bovine serum (EUROBIO) and antibiotics (100 U/ml penicillin and 100 µg/ml streptomycin, Invitrogen) and the appropriate selection antibiotics for stable cell lines (Euromedex, Table 1). For seeding NIH/3T3 cells on coverslips, coverslips were first coated with 20 µg/ml Collagen Type I (MERCK Millipore) and 2 µg/ml fibronectin (Sigma-Aldrich) to increase cell adhesion.

**Table 1:** Stable cell lines.

| Stable cell lines | Selection antibiotics | Reference |
| --- | --- | --- |
| NIH/3T3 GFP-DDB2 | 200 µg/ml Hygromycin | (14) |
| U2OS GFP-DDB2 | 200 µg/ml Hygromycin | This study |
| U2OS H2A.X-SNAP | 100 µg/ml G418 | (15) |
| U2OS H3.1-SNAP | 100 µg/ml G418 | (47) |
| U2OS H3.3-SNAP | 100 µg/ml G418 | (47) |
| RPE-1 H3.1-SNAP | 300 µg/ml G418 | This study |
| RPE-1 H3.3-SNAP | 300 µg/ml G418 | This study |

In order to inhibit DNA replication, we use 10 mM Hydroxyurea (HU, Sigma-Aldrich) and 1 mM cytosine arabinoside (Ara-C, Sigma-Aldrich) in fresh growth medium for 2 h prior to DNA damage induction.

### siRNA and plasmid transfections

siRNA purchased from Eurofins MWG Operon, Sigma Aldrich and Dharmacon (Table 2) were transfected into cells using Lipofectamine RNAiMAX (Invitrogen) following manufacturer’s instructions. The final concentration of siRNA in the culture medium was 50 nM. Cells were harvested 48-72 h after transfection.

**Table 2:**
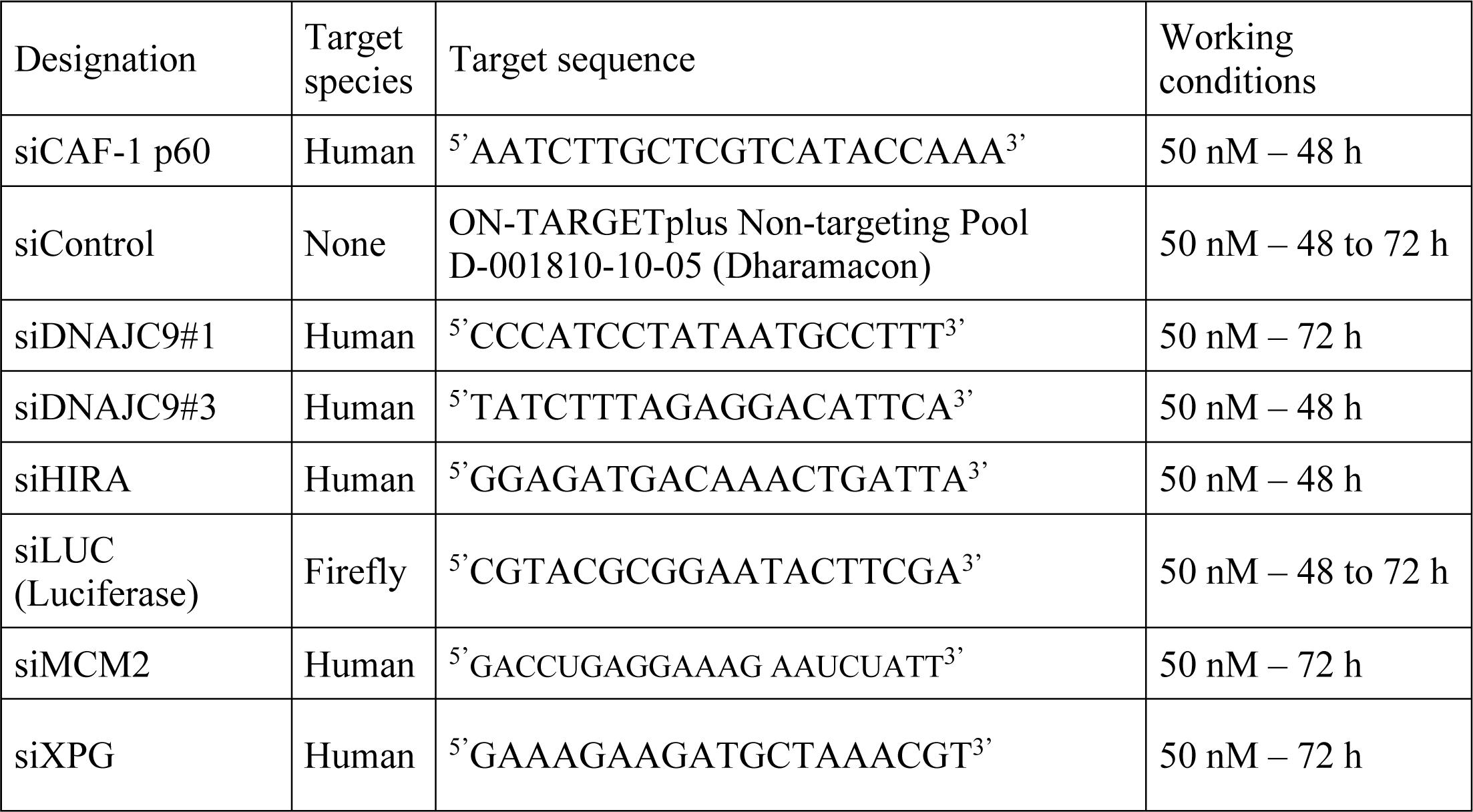
siRNA sequences.

Cells were transfected with plasmid DNA (Table 3) using Lipofectamine 2000 (Invitrogen) according to manufacturer’s instructions. For stable cell line establishment (Table 1), plasmid DNA was transfected into cells at 1 µg/ml final, 48 h before antibiotic selection of clones.

**Table 3:** Plasmids.

| Plasmid | Construct details | Reference |
| --- | --- | --- |
| GFP-DDB2 | Human DDB2 coding sequence (Montpellier Genomic Collections) subcloned into GFP-XPC plasmid replacing XPC | (16) |
| H3.1-SNAP | Human H3C3 coding sequence cloned into pSNAPm (New England Biolabs) | (47) |
| H3.3-SNAP | Human H3F3B coding sequence cloned into pSNAPm (New England Biolabs) | (47) |

### UVC irradiation

Cells grown on glass coverslips (12 mm diameter, thickness No.1.5, Thorlabs) were irradiated with UVC (254 nm) using a low-pressure mercury lamp. Conditions were set using a VLX-3W dosimeter (Vilber Lourmat). For global UVC irradiation, cells in Phosphate Buffer Saline (PBS) were exposed to UVC doses ranging from 4 to 12 J/m^2^ for survival assays and to 10 J/m^2^ in other experiments. For local UVC irradiation (48, 49), cells were covered with a polycarbonate filter (5 μm pore size, Millipore) and irradiated with 150, 300 or 500 J/m^2^ UVC. Irradiated cells were allowed to recover in culture medium for the indicated times before fixation.

For UVC laser micro-irradiation (50), cells were grown on quartz coverslips (25 mm diameter, thickness No.1, SPI supplies) and nuclei were stained by adding Hoechst 33258 (10 μg/mL final, Sigma-Aldrich) to the culture medium 30 min before UVC irradiation. Quartz coverslips were transferred to a Chamlide magnetic chamber (Gataca-systems) on a custom stage insert (Live Cell Instrument) and cells were irradiated for 100 ms using a 2 mW pulsed diode-pumped solid-state laser emitting at 266 nm (RappOptoElectronics, Hamburg GmbH) directly connected to a Zeiss LSM900 confocal microscope adapted for UVC transmission with all-quartz optics. The laser was attenuated using a 0.1% neutral density filter and focused through a 40x/0.6 Ultrafluar glycerol objective with quartz lenses. The laser is controlled by a UGA Firefly module with SysCon2 software (RappOpto). The laser impact has an average size of 1 μm in diameter and damages around 1-2% of the total nuclear volume. The corresponding UVC dose, not directly measurable but estimated by comparing the intensity of the CPD damage generated by the laser and the UVC lamp, is 800 J/m^2^.

### IPOND-R (Isolation of Proteins On Nascent DNA at Repair sites)

#### EdU labeling of repair sites

Cells are seeded in 15-cm dishes to harvest at least 70 million cells per condition on the day of the experiment. In order to label only repair patches, replication is inhibited with 10 mM HU and 1 mM Ara-C (Sigma-Aldrich) in fresh growth medium for 2 h at 37°C prior to global UVC irradiation (10 J/m^2^) in PBS supplemented with 10 mM HU and 1 mM Ara-C to maintain the inhibition of replication. A non-irradiated sample is used as negative control. After irradiation, cells are incubated in fresh medium containing the replication inhibitor cocktail and 10 µM Ethynyl-deoxyUridine (EdU, Euromedex) for 1 h at 37°C. At this step, a chase can be performed in fresh medium supplemented with 10 µM Thymidine (Sigma-Aldrich).

#### Cell fixation and permeabilization

Cells are fixed with 1% Formaldehyde (Sigma-Aldrich) for 15 minutes on a rocking platform. The fixation is stopped by adding 1.33 M Glycine (Sigma-Aldrich) for 5 minutes on the rocking platform. Cells are washed twice in cold PBS, scrapped in 1% BSA solution in PBS and pelleted for 5 min at 300g at 4°C. Cell pellets are permeabilized with 1% Triton X-100 (1 mL/ 10 million cells, Euromedex) for 30 min at room temperature and washed twice with cold 1% BSA solution.

#### EdU biotinylation by Click-it chemistry

Cell pellets are resuspended in a Click-it reaction cocktail containing 2 mM CuSO_4_ (Sigma-Aldrich), 1 mM Tris((1-hydroxy-propyl-1H-1,2,3-triazol-4-yl)methyl)amine (THPTA, Euromedex), 10 mM Sodium Ascorbate (Sigma-Aldrich), 5 mM Amino-guanidine hydrochloride (Sigma-Aldrich), and 10 μM Biotin Picolyl azide or Biotin azide (Sigma-Aldrich). Cell suspensions are left on a rotating wheel at room temperature for 1 h. Cells are then washed twice in cold 1% BSA solution containing 0.5X EDTA-free Protease inhibitor cocktail (PIC, Roche), and once in cold PBS. A negative control without biotin reagent was performed to check for the specificity of the Streptavidin pull-down.

#### Cell lysis and sonication

Cell pellets are resuspended in lysis buffer (200 ul per 15 million cells) containing 1% SDS, 50 mM Tris pH 7.5, and 1X PIC. The cell suspensions are transferred to 1.5 ml sonication tubes (Diagenode) and sonication is performed using a Bioruptor Pico (Diagenode) with 5-10 cycles of 30 sec ON and 30 sec OFF. The sonicated samples are transferred to new Eppendorf tubes and centrifuged 10 min at 16,000 g at 4°C. The supernatants are transferred to Low Protein Binding tubes (Thermo Fisher Scientific).

#### Analysis of DNA shearing

A 20 μl aliquot of each cell lysate is incubated with 200 mM NaCl and 200 μg/ml RNAse A (Millipore) overnight at 65°C. After addition of 100 μg/ml Proteinase K (Sigma-Aldrich), samples are incubated for a further 2 h at 45°C prior to DNA purification using the PCR purification kit from Macherey-Naegel following manufacturer’s instructions. DNA samples are analyzed with a TapeStation (Agilent) to check that the average fragment size is about 500 bp.

#### Streptavidin capture of biotin-labeled nascent DNA and associated proteins

The remaining solution is resuspended in an equal volume of cold PBS supplemented with 1 X PIC to dilute the SDS present in the sonication buffer. At this stage, input samples are collected, diluted in Laemmli buffer (50 mM Tris-HCl pH 6.8, 1.6% Sodium Dodecyl Sulfate (SDS), 8% glycerol, 4% β-mercaptoethanol, 0.0025% bromophenol blue) and decrosslinked by heating for 5 min at 95°C.

For streptavidin capture, Dynabeads MyOne Streptavidin-C1 beads (Invitrogen) are washed in lysis buffer and in PBS supplemented with 1X PIC and then are added to cell lysates (1 µl beads per 1 million cells) and incubated overnight at 4°C on a rotating wheel. After capture, beads are washed once in lysis buffer (1 ml wash solution per 10 million cells), once in 1M NaCl supplemented with 1X PIC, followed by 3 additional washes in lysis buffer.

For western blot analysis, the beads are resuspended in Laemmli buffer and decrosslinking is achieved by heating for 5 min at 95°C.

For mass spectrometry analysis, beads are transferred to new Low Protein binding tubes and washed in 50 mM Tris-HCl pH 8 before resuspension in Laemmli buffer and decrosslinking by heating for 5 min at 95°C. Ten percent of the protein solution is kept for a Silver staining analysis.

### Mass spectrometry-based proteomic analyses

Proteins from three and four IPOND-R biological replicates, purified 1h and 24h after UV irradiation, respectively (-UV, negative control) were solubilized in Laemmli buffer (50 mM Tris-HCl pH 6.8, 1.6% Sodium Dodecyl Sulfate (SDS), 8% glycerol, 4% β-mercaptoethanol, 0.0025% bromophenol blue) and separated by SDS-PAGE (Mini Protean TGX, 4-20%, Biorad), before staining with Coomassie blue R-250 (Bio Basic Canada Inc.). The band corresponding to streptavidin was discarded and the remaining proteins were digested in-gel using modified trypsin (Promega, sequencing grade) as previously described (51). The resulting peptides were analyzed by online nanoliquid chromatography coupled to MS/MS (Ultimate 3000 RSLCnano and Q-Exactive HF, Thermo Fisher Scientific, for the 1h dataset, and Ultimate 3000 RSLCnano and Orbitrap Exploris 480, Thermo Fisher Scientific, for the 24h dataset) using a 120-min gradient. For this purpose, the peptides were sampled on a precolumn (300 μm x 5 mm PepMap C18, Thermo Scientific) and separated in a 75 μm x 250 mm C18 column (Reprosil-Pur 120 C18-AQ, 1.9 μm, Dr. Maisch, for the 1h dataset, and Aurora Generation 2, 1.6 µm, IonOpticks, for the 24h dataset). The MS and MS/MS data were acquired using Xcalibur (Thermo Fisher Scientific).

Peptides and proteins were identified by Mascot (version 2.8.0, Matrix Science) through concomitant searches against the Uniprot database (20230719 download, 207’981 sequences) and a homemade database containing the sequences of classical contaminant proteins found in proteomic analyses (human keratins, trypsin…, 250 sequences). Trypsin/P was chosen as the enzyme and two missed cleavages were allowed. Precursor and fragment mass error tolerances were set at respectively at 10 and 20 ppm. Peptide modifications allowed during the search were: Carbamidomethyl (C, fixed), Acetyl (Protein N-term, variable) and Oxidation (M, variable). The Proline software (52) version 2.2.0 was used for the compilation, grouping, and filtering of the results (conservation of rank 1 peptides, peptide length ≥ 6 amino acids, false discovery rate of peptide-spectrum-match identifications < 1% (53), and minimum of one specific peptide per identified protein group). MS data have been deposited to the ProteomeXchange Consortium via the PRIDE partner repository (54) with the dataset identifier PXD055129. Proline was then used to perform a MS1 label-free quantification of the identified protein groups based on razor and specific peptides.

Statistical analysis was performed using the ProStaR software(55) based on the quantitative data obtained with the different biological replicates analyzed for each condition. Proteins identified in the contaminant database, proteins identified by MS/MS in less than two replicates of one condition, and proteins quantified in less than all replicates of one condition were discarded. After log2 transformation, abundance values were normalized using the variance stabilizing normalization (vsn) method applied condition-wise, before missing value imputation (SLSA algorithm for partially observed values in the condition and DetQuantile algorithm for totally absent values in the condition). Statistical testing was conducted with limma, whereby differentially expressed proteins were selected using a log2 (Fold Change) cut-off of 1 and a p-value cut-off of 0.01, allowing to reach a false discovery rate inferior to 3% according to the Benjamini-Hochberg estimator. Proteins found differentially abundant but identified by MS/MS in less than two replicates, and detected in less than all replicates, in the condition in which they were found to be more abundant were invalidated (p-value = 1).

### Data visualization and Gene ontology analysis

Biological process gene ontology terms were extracted from the PANTHER database (v.18 2023-08-01 release) and analyzed through the PANTHER Overrepresentation Test (released 2023-07-05). Significance was calculated using a Fisher test with a false-discovery rate correction. The list of EpiFactors was extracted from https://epifactors.autosome.ru/public_dataversion2.0 (56, 57) curated based on most recent literature. The proteins detected by IPOND-R coupled to mass spectrometry were visualized using Volcano plots generated with R studio 2021.09.0 using ggplot2 version 3.3.5, ggrepel version 0.9.1, and dplyr version 1.1.2. Venn diagrams were generated with eulerr version 6.1. Statistical analyses and all other graphs were generated using GraphPad Prism version 8.4.3, and test details and p-values are referred to in the figure legends.

### EdU-labeling of replicating cells and repair sites

To visualize replication foci, 10 µM Ethynyl-deoxyUridine (EdU) was incorporated into cells on glass coverslips during 15 min at 37°C and revealed using the Click-It EdU Alexa Fluor 488 or 594 Imaging kit (Invitrogen) according to manufacturer’s instructions. To localize the sites of UV damage repair, cells were incubated with 10 µM EdU for 1h30 after local UVC irradiation and EdU was revealed using the Click-It EdU Alexa Fluor 488, 594 or 647 Imaging kit (Invitrogen).

### Immunofluorescence

Cells grown on coverslips were either fixed directly with 2% paraformaldehyde (Electron Microscopy Sciences) for 10 min and permeabilised for 5 min with 0.5% Triton X-100 in PBS or cells were pre-extracted before fixation with 0.5% Triton X-100 in CSK buffer (Cytoskeletal buffer: 10 mM PIPES pH 7.0, 100 mM NaCl, 300 mM sucrose, 3 mM MgCl_2_) for 5 min at room temperature to remove soluble proteins. For the detection of UVC photoproducts, the DNA was denatured with 2N HCl for 10 min at 37°C (6,4-PP detection) or with 0.5 M NaOH for 5 min at room temperature (CPD detection). Since this denaturation quenches GFP fluorescence, when CPD detection was combined with the visualization of GFP-DDB2, immunofluorescence was performed in two steps starting with GFP immunodetection using a rat anti-GFP antibody (Table 4) followed by fixation, denaturation and CPD immunodetection. Samples were blocked for 10 min in 5% BSA (Bovine Serum Albumin, Sigma-Aldrich) in PBT (PBS 0.1% Tween-20), followed by 45 min incubation with primary antibodies and 30 min incubation with secondary antibodies coupled to AlexaFluor dyes (Table 4) diluted in blocking buffer. Coverslips were mounted in Vectashield medium with DAPI (Vector laboratories).

**Table 4:**
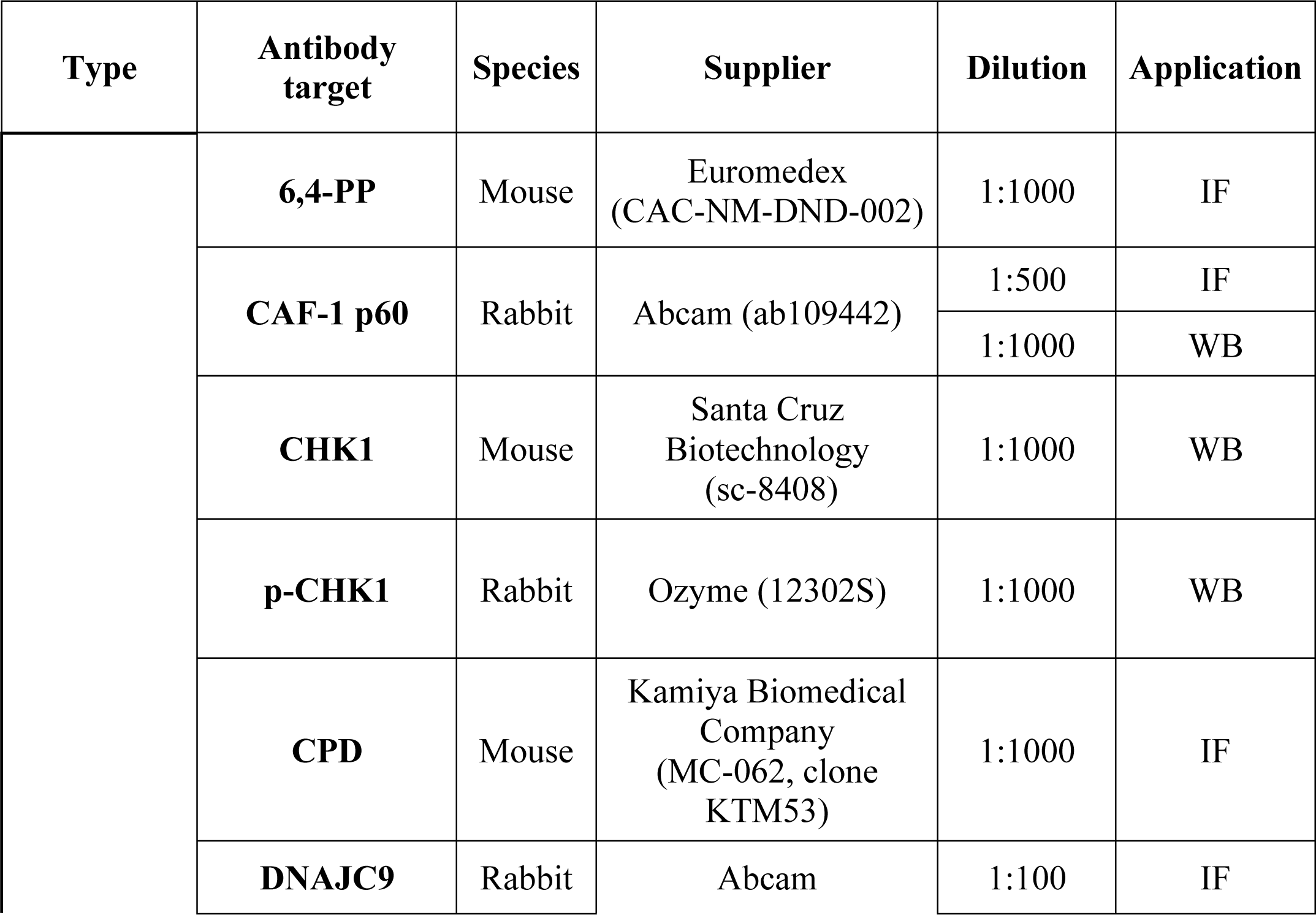
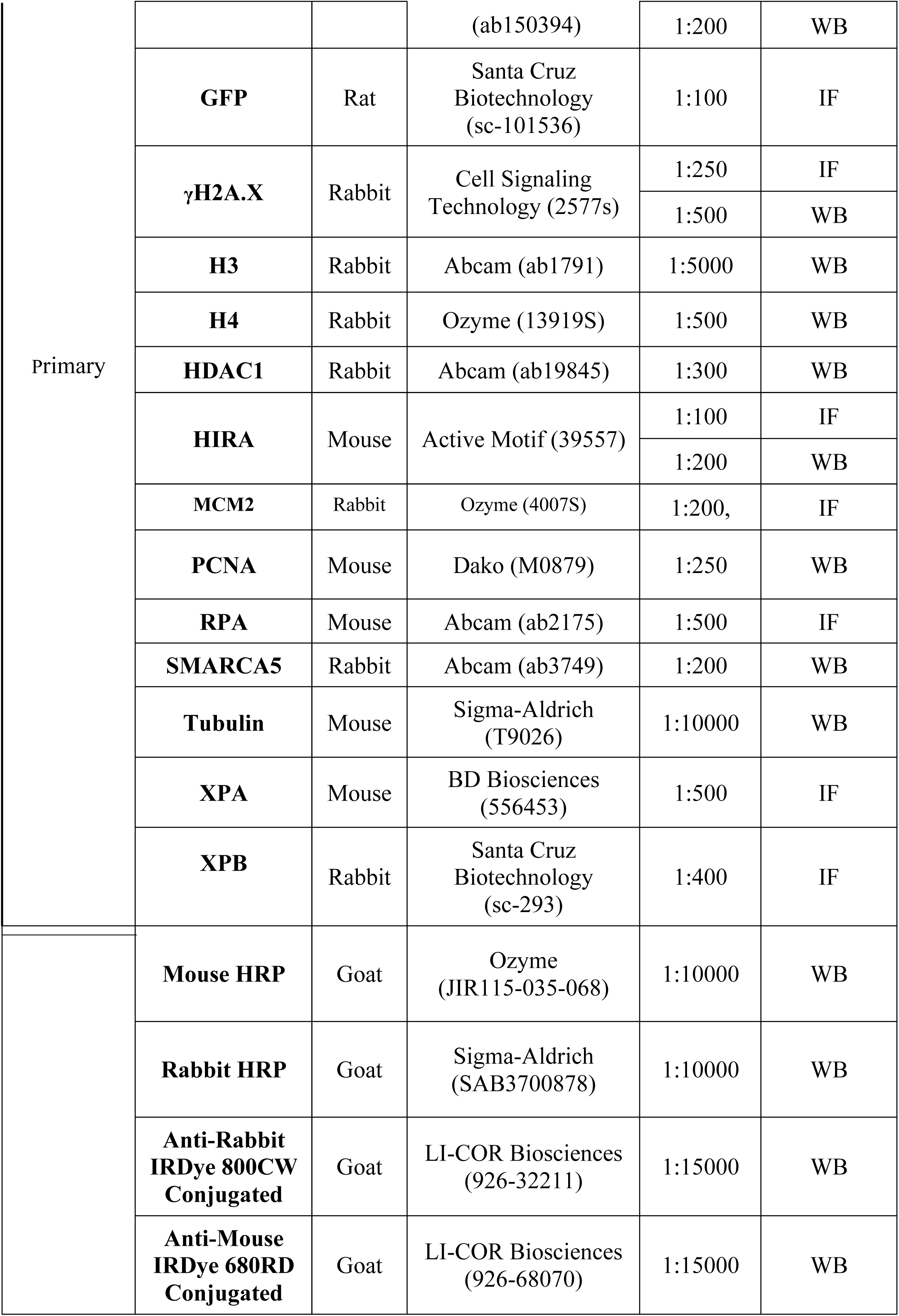
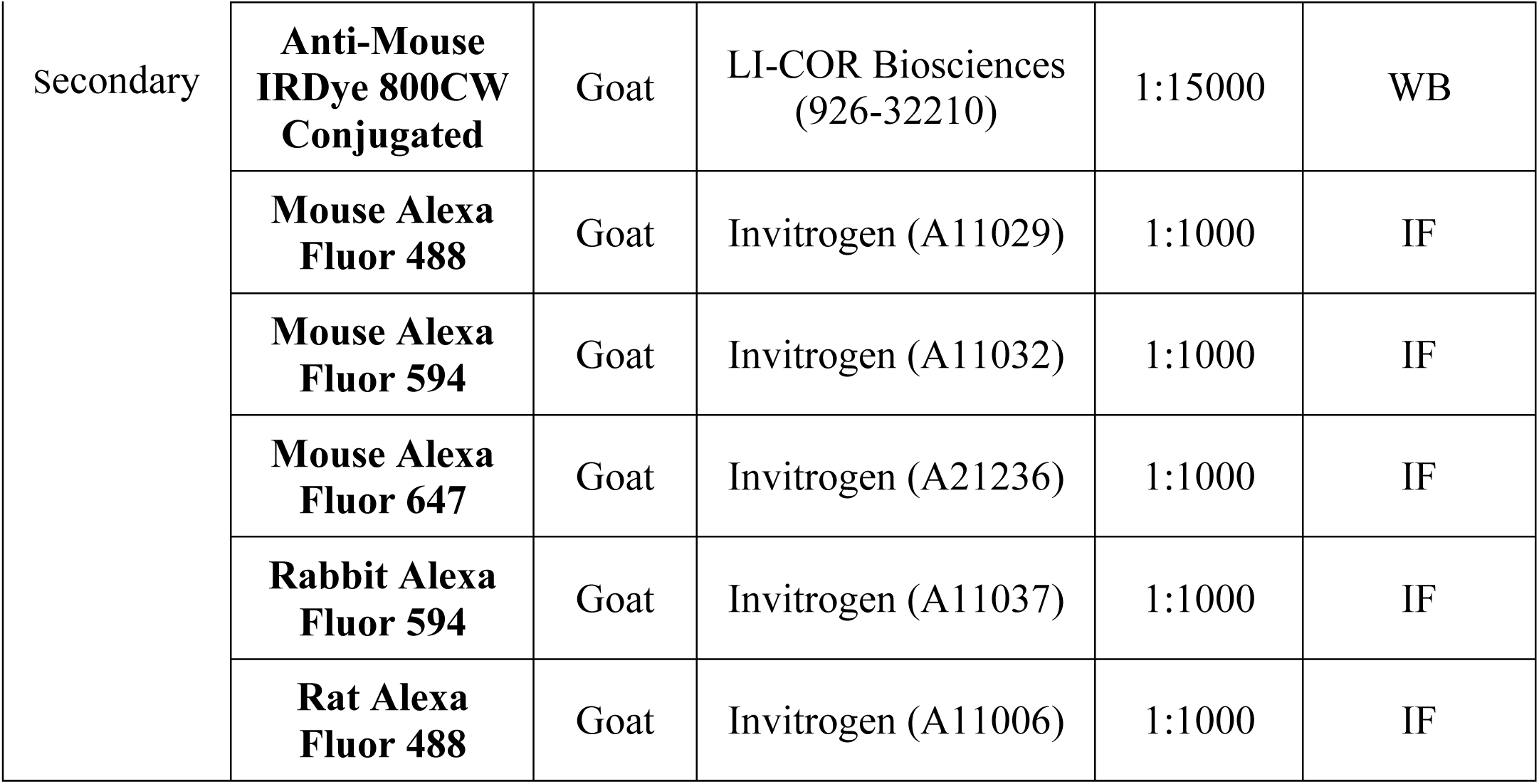
Antibodies.

### SNAP-tag labelling of histones

For specific labelling of newly synthesized histones (33, 58), cells were grown on glass coverslips and pre-existing SNAP-tagged histones were first quenched by incubating cells with 10 μM of the non-fluorescent substrate SNAP-cell Block (New England Biolabs) for 30 min followed by a 30 min-wash in fresh medium and a 2 h (U2OS) or 4 h (RPE-1) chase. The new SNAP-tagged histones synthesized during the chase were fluorescently labelled with 2 μM of the red-fluorescent reagent SNAP-cell TMR star (New England Biolabs) during a 15 min-pulse step followed by 30 min wash in fresh medium. Cells were subsequently permeabilised with Triton X-100, fixed and processed for immunostaining. Cells were irradiated with a UVC lamp before the pulse step.

For specific labelling old histones, cells were grown on glass coverslips and pre-existing SNAP-tagged histones were fluorescently labelled with 2 μM of the red-fluorescent reagent SNAP-cell TMR star (New England Biolabs) during a 30 min-pulse step followed by 30 min wash in fresh medium. Cells were irradiated with a UVC lamp 48 h after the pulse step. Cells were subsequently permeabilised with Triton X-100, fixed and processed for immunostaining.

For labelling all histones, cells were grown on glass coverslips and SNAP-tagged histones were fluorescently labelled with 2 μM of the red-fluorescent reagent SNAP-cell TMR star (New England Biolabs) during a 30 min-pulse step. Cells were subsequently permeabilised with Triton X-100, fixed and processed for immunostaining. Cells were irradiated with a UVC lamp before the pulse step.

### Image acquisition and analysis

Fluorescence imaging was performed with a Leica DMI6000 epifluorescence microscope using a Plan-Apochromat 40x/1.3 oil objective. Images were captured using a CCD camera (Photometrics) and Metamorph software. Images were assembled with Adobe Photoshop. Live cell imaging coupled to UVC laser micro-irradiation was performed using a 40x/0.6 Ultrafluar Glycerol objective on a Zeiss LSM900 confocal microscope. Images were captured using Zen blue software, and analysed with ImageJ (U. S. National Institutes of Health, Bethesda, Maryland, USA, http://imagej.nih.gov/ij/). Nuclei were segmented based on DAPI or Hoechst staining, and UVC-damaged regions based on GFP-DDB2 fluorescence or immunostaining for UV damage or repair factors using custom-made ImageJ macros.

### Cell extracts and western blot

Total extracts were obtained by scraping cells on plates or resuspending cell pellets in Laemmli buffer (50 mM Tris-HCl pH 6.8, 1.6% Sodium Dodecyl Sulfate (SDS), 8% glycerol, 4% β-mercaptoethanol, 0.0025% bromophenol blue) followed by 5 min denaturation at 95°C.

For western blot analysis, extracts were run on 4%–20% Mini-PROTEAN TGX gels (Bio-Rad) in running buffer (200 mM glycine, 25 mM Tris, 0.1% SDS). Proteins were transferred onto nitrocellulose membranes (0.2 μm, Amersham Protran) for 30 min at 15V with a Trans-Blot SD semidry transfer cell (Bio-Rad) in transfer buffer (50 mM Tris, 40 mM glycine, 0.04% SDS, 15% ethanol). Total proteins were revealed by Pierce® Reversible Stain (Thermo Scientific). Proteins of interest were probed using the appropriate primary and Horse Radish Peroxidase (HRP)-conjugated secondary antibodies (Table 4), detected using SuperSignal West Pico or Femto chemiluminescence substrates (Pierce) on hyperfilms MP (Amersham). When fluorescence detection was used instead of chemi-luminescence, secondary antibodies were conjugated to IRDye 680RD or 800CW (Table 4), membranes were scanned with an Odyssey Fc-imager (LI-COR Biosciences) and analysed with Image Studio Lite software using total protein stain for normalization.

### Flow cytometry

For cell cycle analysis, cells were fixed in ice-cold 70% ethanol before DNA staining with 50 ug/ml propidium iodide (Sigma-Aldrich) in PBS containing 0.05% Tween and 0.5 mg/ml RNase A (USB/Affymetrix). DNA content was analysed by flow cytometry using a BD FACScalibur flow cytometer (BD Biosciences) and FlowJo software (TreeStar).

### Colony forming assays

Cells were replated 48 h after siRNA transfection and exposed to global UVC irradiation (4, 8 and 12 J/m^2^) the following day. Colonies were stained 12 days later with 0.5% crystal violet/20% ethanol and counted. Results were normalized to plating efficiencies.

### Statistical analyses

Percentages of positively stained cells were obtained by scoring at least 150 cells in each experiment. Statistical tests were performed using GraphPad Prism. P-values for mean comparisons between two groups were calculated with a Student’s t-test with Welch’s correction when necessary. Multiple comparisons were performed by one-way or two-way ANOVA. Comparisons of clonogenic survival were based on non-linear regression with a polynomial quadratic model. ns: non-significant, *: p < 0.05, **: p < 0.01, ***: p < 0.001.

## ACKNOWLEDGMENTS

We thank the EPI^2^ Imaging platform and in particular Sandra Piquet - UMR7216 Epigenetics and Cell Fate Centre for access to instruments and technical advice.

This work was supported by the European Research Council (ERC-2018-CoG-818625 “REMIND”). The proteomic experiments were partially supported by Agence Nationale de la Recherche under projects ProFI (Proteomics French Infrastructure, ANR-10-INBS-08) and GRAL, a program from the Chemistry Biology Health (CBH) Graduate School of University Grenoble Alpes (ANR-17-EURE-0003).

## AUTHOR CONTRIBUTIONS

A.P., A.C., E.P. and S.E.P. designed and performed experiments, analysed the data and wrote the manuscript. J.N.D and Y.C. performed MS-based proteomic analyses. S.E.P. supervised the project.

---

**Supplementary figure 1: Effect of transient replication inhibition on DNA repair and chromatin dynamics**

(**A-B**) Immunodetection of repair-associated factors RPA (A), γH2A.X (B) at sites of local UVC irradiation (marked by CPD or XPB) in U2OS cells treated or not with replication inhibitors (Replic. inhib.).

(**C**) Immunoblot for the DNA damage checkpoint marker phospho-CHK1 (p-CHK1) and its non-activated form (CHK1) in total extracts from U2OS cells treated or not with replication inhibitors (Replic. inhib.) and harvested 1 h or 24 h post UVC. The corresponding quantifications are shown below (mean ± SD from 3 independent experiments).

(**D**) Cell cycle distribution of U2OS cells treated or not with replication inhibitors (Replic. inhib.) and harvested 1 h or 24 h post UVC irradiation (mean ± SD from 3 independent experiments).

(**E**) Immunodetection of the repair factor XPB at sites of local UVC irradiation (marked by CPD) in U2OS cells treated or not with replication inhibitors (Replic. inhib.).

(**F-G)** Time course of UV lesion removal: CPD (F) and 6,4-PP (G) mean nuclear intensities are measured by immunofluorescence after global UVC irradiation in U2OS cells treated or not with replication inhibitors (Replic. inhib.). The data are normalized to T0 (time of irradiation). Mean ± SEM from 3 independent experiments.

(**H, J**) Detection and quantification of newly synthesized H3.1-SNAP (H) and H3.3-SNAP (J) histones at sites of UVC irradiation (marked by CPD and XPB, respectively) in U2OS cells stably expressing SNAP-tagged H3 variants and treated or not with replication inhibitors (Replic. inhib.). New H3.1-SNAP signal at damage sites is quantified in non-S phase cells by excluding from the analysis nuclei showing pan-nuclear EdU staining.

(**I, K**) Immunodetection of CAF-1 (I) and HIRA (K) chaperones at sites of UVC irradiation (marked by CPD and XPB, respectively) in U2OS cells treated or not with replication inhibitors (Replic. inhib.). CAF-1 signal at damage sites is quantified in non-S phase cells by excluding from the analysis nuclei showing pan-nuclear CAF-1 staining.

Dot plots: mean ± SD from at least 56 cells scored in 3 independent experiments (the mean of each experiment is shown as a grey triangle). Statistical significance is calculated with one-way ANOVA (A-C), two-way ANOVA (D), a two-sided Student’s t-test with Welch’s correction (E, H-K), or non-linear regression with a polynomial quadratic model (F-G). Scale bars, 10 μm.

**Supplementary figure 2: EdU-dependent labeling of repair synthesis**

**(A)** Detection and quantification of EdU nuclear signal by Click-it chemistry performed with biotin azide or biotin picolyl azide 1 h after global UV irradiation in U2OS cells pre-treated for 2 h with replication inhibitors. Biotinylated EdU was revealed with fluorescently-labelled streptavidin. Mean ± SD from at least 43 cells scored in 3 independent experiments.

**(B)** Persistance of the biotinylated EdU signal over time after global UV irradiation in U2OS cells pre-treated for 2 h with replication inhibitors. EdU was biotinylated using biotin picolyl azide and revealed as in (A). Mean ± SD from at least 79 cells scored in 3 independent experiments.

**(C)** Scheme of the protocol used to compare EdU incorporation in euchromatin vs. heterochromatin after UVC damage in NIH/3T3 murine fibroblasts, which display DAPI-dense chromocenters (constitutive heterochromatin). The EdU pulse was performed 0 h or 3 h post UVC irradiation to capture early and later repair events. Representative immunostaining and quantification of EdU enrichment in heterochromatin/euchromatin in the two experimental conditions (mean ± SD from at least 50 cells scored in 3 independent experiments). Statistical significance is calculated with a two-sided Student’s t-test with Welch’s correction (A, C) and with a multiple t-test (B). Scale bars, 10 μm.

**Supplementary figure 3: Factors enriched at sites of UV damage repair**

**(A)** Venn diagram showing proteins identified by IPOND-R at 1 h post UVC irradiation in U2OS cells (this study, +/-UV fold-change (FC)>= 2 and limma p-value<= 0.01), proteins from the aniFOUND dataset 4 h post UVC in human skin fibroblasts (Stefos et al., group A, FDR<0.1), and proteins from the chromatin proteome 3 h post UV in HEK293 cells (Boeing et al., z-score>1).

**(B)** Top 25 biological process terms in IPOND-R 1 h-specific hits (Panther DB gene ontology analysis). The False Discovery Rate is represented on the x-axis, and the log2-fold enrichment of the biological processes by the circle size.

**Supplementary figure 4: DNAJC9 is dispensable for the repair of UV-induced DNA damage**

**(A)** Immunodetection of DNAJC9 in U2OS cells treated with the indicated siRNAs (siLuc, control). The DNAJC9 signal in the nucleus is normalized to the mean of the corresponding siLuc experiment.

**(B)** Immunodetection of XPB at damage sites (marked by CPD) 20 min post local UVC irradiation in U2OS cells treated with the indicated siRNAs (siLuc, negative control). The dot plot shows the mean signal intensity of XPB at damage sites normalized to the mean of the corresponding siLuc experiment.

**(C)** EdU detection by Click-it chemistry at damage sites (marked by CPD) 1 h post local UVC irradiation in U2OS cells treated with the indicated siRNAs (siLuc, negative control). The dot plot shows the the EdU signal at damage sites outside S-phase normalized to the mean of the corresponding siLuc experiment.

**(D)** Time course of UV lesion removal: CPD (left) and 6,4-PP (middle) mean nuclear intensities are measured by immunofluorescence after global UVC irradiation in U2OS cells treated with the indicated siRNAs (siLuc, negative control; siXPG, positive control). The data are normalized to the siLuc condition at T0 (time of irradiation). Mean ± SEM from 3-4 independent experiments. Statistical significance is given by non-linear regression with a polynomial quadratic model. Control of DNAJC9 depletion by immunofluorescence in U2OS cells is shown on the right.

**(E)** Clonogenic survival to UVC irradiation in U2OS cells treated with the indicated siRNAs (siLuc, negative control; siXPG, positive control). Statistical significance is given by non-linear regression with a polynomial quadratic model. The efficiency of DNAJC9 depletion is controlled by western blot (right panel).

Dot plots: mean ± SD from at least 259 cells scored in 3-4 independent experiments (the mean of each experiment is represented by a grey triangle). Statistical significance is calculated with a two-sided Student’s t-test with Welch’s correction. Scale bars, 10 μm.

**Supplementary figure 5: DNAJC9 stimulates the deposition of newly synthesized H3 variants**

(**A-B**) Detection and quantification of newly synthesized H3.1-SNAP (A) and H3.3-SNAP (B) histones at sites of UVC irradiation (marked by CAF-1 p60 and XPA, respectively) in U2OS cells stably expressing SNAP-tagged H3 variants and treated with the indicated siRNAs (siLuc, control). The incorporation of new H3.1-SNAP histones is quantified in cells outside S-phase. The percentages of signal decrease observed in siDNAJC9-treated cells are indicated on the right. The efficiency of DNAJC9 depletion and the impact on CAF-1 and HIRA chaperone levels are monitored by western blot (Tubulin and H3, loading controls).

(**C-D**) Quantification of new H3.1-SNAP (C, left panel) and H3.3-SNAP (D, left panel) incorporation at sites of UVC irradiation in RPE-1 cells stably expressing SNAP-tagged H3 variants and treated with the indicated siRNAs (siLuc, control). The percentages of signal decrease observed in DNAJC9-depleted cells are indicated. The efficiency of DNAJC9 depletion is verified by immunofluorescence and quantification of nuclear levels (right panels).

**(E)** Quantification of new H2A.X-SNAP incorporation at sites of UVC irradiation (marked by CPD) in U2OS cells stably expressing H2A.X-SNAP and treated with the indicated siRNAs (siLuc, negative control; siXPG, positive control impairing new H2A.X deposition at UV sites).

**(F)** Percentage of cells in S-phase determined by CAF-1 p60 immunostaining of replication foci in U2OS cells treated with the indicated siRNAs (siLuc, control). Mean ± SD from at least 40 cells scored in 3 independent experiments.

(**G-H**) Quantification of new H3.1-SNAP (G) and new H3.3-SNAP (H) incorporation in chromatin in undamaged U2OS cells stably expressing SNAP-tagged H3 variants and treated with the indicated siRNAs (siLuc, control). The incorporation of new H3.1-SNAP histones is quantified in S-phase cells based on CAF-1 p60 immunostaining.

(**I-L**) Efficiency of DNAJC9 (I, K), CAF-1 p60 (J) and HIRA (L) depletion controlled by immunofluorescence in U2OS cells stably expressing SNAP-tagged H3 variants and treated with the indicated siRNAs (siLuc, control).

Dot plots: mean ± SD from at least 69 cells scored in 3-4 independent experiments (the mean of each experiment is represented by a grey triangle). Statistical significance is calculated by one-way ANOVA with multiple comparison (E, I-L) or with a two-sided Student’s t-test with Welch’s correction (A-D, F-H). Scale bars, 10 μm.

**Supplementary figure 6: MCM2 cooperates with DNAJC9 in controlling histone dynamics at sites of UV damage repair**

(**A-B**) Efficiency of DNAJC9 and MCM2 depletion controlled by immunofluorescence in U2OS cells stably expressing SNAP-tagged H3 variants and treated with the indicated siRNAs (siLuc, control). Statistical significance is calculated by one-way ANOVA with multiple comparison.

## REFERENCES

1. Allis, C.D. and Jenuwein, T. (2016) The molecular hallmarks of epigenetic control. Nat. Rev. Genet., 17, 487–500.

2. Millán-Zambrano, G., Burton, A., Bannister, A.J. and Schneider, R. (2022) Histone post-translational modifications — cause and consequence of genome function. Nat. Rev. Genet., 23, 563–580.

3. Greenberg, M.V.C. and Bourc’his, D. (2019) The diverse roles of DNA methylation in mammalian development and disease. Nat. Rev. Mol. Cell Biol., 20, 590–607.

4. Martire, S. and Banaszynski, L.A. (2020) The roles of histone variants in fine-tuning chromatin organization and function. Nat. Rev. Mol. Cell Biol., 21, 522–541.

5. Hammond, C.M., Strømme, C.B., Huang, H., Patel, D.J. and Groth, A. (2017) Histone chaperone networks shaping chromatin function. Nat. Rev. Mol. Cell Biol., 18, 141–158.

6. Clapier, C.R., Iwasa, J., Cairns, B.R. and Peterson, C.L. (2017) Mechanisms of action and regulation of ATP-dependent chromatin-remodelling complexes. Nat. Rev. Mol. Cell Biol., 18, 407–422.

7. Bannister, A.J. and Kouzarides, T. (2011) Regulation of chromatin by histone modifications. Cell Res, 21, 381–395.

8. Flavahan, W.A., Gaskell, E. and Bernstein, B.E. (2017) Epigenetic plasticity and the hallmarks of cancer. Science, 357.

9. Brookes, E. and Shi, Y. (2014) Diverse Epigenetic Mechanisms of Human Disease. Annu. Rev. Genet., 48, 1–32.

10. Ferrand, J., Plessier, A. and Polo, S.E. (2021) Control of the chromatin response to DNA damage: Histone proteins pull the strings. Semin. Cell Dev. Biol., 113, 75–87.

11. Stewart-Morgan, K.R., Petryk, N. and Groth, A. (2020) Chromatin replication and epigenetic cell memory. Nat. Cell Biol., 22, 361–371.

12. Hoeijmakers, J.H.J. (2009) DNA Damage, Aging, and Cancer. N. Engl. J. Med., 361, 1475–1485.

13. Dabin, J., Mori, M. and Polo, S.E. (2023) The DNA damage response in the chromatin context: A coordinated process. Curr. Opin. Cell Biol., 82, 102176.

14. Fortuny, A., Chansard, A., Caron, P., Chevallier, O., Leroy, O., Renaud, O. and Polo, S.E. (2021) Imaging the response to DNA damage in heterochromatin domains reveals core principles of heterochromatin maintenance. Nat. Commun., 12, 2428.

15. Piquet, S., Parc, F.L., Bai, S.-K., Chevallier, O., Adam, S. and Polo, S.E. (2018) The Histone Chaperone FACT Coordinates H2A.X-Dependent Signaling and Repair of DNA Damage. Mol. Cell, 72, 888–901.e7.

16. Adam, S., Dabin, J., Chevallier, O., Leroy, O., Baldeyron, C., Corpet, A., Lomonte, P., Renaud, O., Almouzni, G. and Polo, S.E. (2016) Real-Time Tracking of Parental Histones Reveals Their Contribution to Chromatin Integrity Following DNA Damage. Mol. Cell, 64, 65–78.

17. Adam, S., Polo, S.E. and Almouzni, G. (2013) Transcription Recovery after DNA Damage Requires Chromatin Priming by the H3.3 Histone Chaperone HIRA. Cell, 155, 963.

18. Polo, S.E., Roche, D. and Almouzni, G. (2006) New Histone Incorporation Marks Sites of UV Repair in Human Cells. Cell, 127, 481–493.

19. Dinant, C., Ampatziadis-Michailidis, G., Lans, H., Tresini, M., Lagarou, A., Grosbart, M., Theil, A.F., Cappellen, W.A. van Kimura, H., Bartek, J., et al. (2013) Enhanced Chromatin Dynamics by FACT Promotes Transcriptional Restart after UV-Induced DNA Damage. Mol. Cell, 51, 469–479.

20. Sirbu, B.M., McDonald, W.H., Dungrawala, H., Badu-Nkansah, A., Kavanaugh, G.M., Chen, Y., Tabb, D.L. and Cortez, D. (2013) Identification of Proteins at Active, Stalled, and Collapsed Replication Forks Using Isolation of Proteins on Nascent DNA (iPOND) Coupled with Mass Spectrometry*. J. Biol. Chem., 288, 31458–31467.

21. Dungrawala, H. and Cortez, D. (2014) Purification of proteins on newly synthesized DNA using iPOND. *Methods Mol. Biol. (Clifton*, NJ), 1228, 123–31.

22. Zeeland, A.A. van Smith, C.A. and Hanawalt, P.C. (1981) Sensitive determination of pyrimidine dimers in DNA of UV-irradiated mammalian cells Introduction of T4 endonuclease V into frozen and thawed cells. Mutat. Res.Fundam. Mol. Mech. Mutagen., 82, 173–189.

23. Wang, L., Cao, X., Yang, Y., Kose, C., Kawara, H., Lindsey-Boltz, L.A., Selby, C.P. and Sancar, A. (2022) Nucleotide excision repair removes thymidine analog 5-ethynyl-2′-deoxyuridine from the mammalian genome. Proc. Natl. Acad. Sci. United States Am., 119, e2210176119.

24. Hu, J., Adebali, O., Adar, S. and Sancar, A. (2017) Dynamic maps of UV damage formation and repair for the human genome. Proc. Natl. Acad. Sci., 114, 6758–6763.

25. Adar, S., Hu, J., Lieb, J.D. and Sancar, A. (2016) Genome-wide kinetics of DNA excision repair in relation to chromatin state and mutagenesis. Proc. Natl. Acad. Sci., 113, E2124– E2133.

26. Stefos, G.C., Szantai, E., Konstantopoulos, D., Samiotaki, M. and Fousteri, M. (2021) aniFOUND: analysing the associated proteome and genomic landscape of the repaired nascent non-replicative chromatin. Nucleic Acids Res., 49, gkab144-.

27. Boeing, S., Williamson, L., Encheva, V., Gori, I., Saunders, R.E., Instrell, R., Aygün, O., Rodriguez-Martinez, M., Weems, J.C., Kelly, G.P., et al. (2016) Multiomic Analysis of the UV-Induced DNA Damage Response. Cell Rep., 15, 1597–1610.

28. Huang, H., Strømme, C.B., Saredi, G., Hödl, M., Strandsby, A., González-Aguilera, C., Chen, S., Groth, A. and Patel, D.J. (2015) A unique binding mode enables MCM2 to chaperone histones H3–H4 at replication forks. Nat. Struct. Mol. Biol., 22, 618–626.

29. Richet, N., Liu, D., Legrand, P., Velours, C., Corpet, A., Gaubert, A., Bakail, M., Moal-Raisin, G., Guerois, R., Compper, C., et al. (2015) Structural insight into how the human helicase subunit MCM2 may act as a histone chaperone together with ASF1 at the replication fork. Nucleic Acids Res., 43, 1905–1917.

30. Han, C., Chen, T., Li, N., Yang, M., Wan, T. and Cao, X. (2007) HDJC9, a novel human type C DnaJ/HSP40 member interacts with and cochaperones HSP70 through the J domain. Biochem. Biophys. Res. Commun., 353, 280–285.

31. Piette, B.L., Alerasool, N., Lin, Z.-Y., Lacoste, J., Lam, M.H.Y., Qian, W.W., Tran, S., Larsen, B., Campos, E., Peng, J., et al. (2021) Comprehensive interactome profiling of the human Hsp70 network highlights functional differentiation of J domains. Mol. Cell, 81, 2549–2565.e8.

32. Hammond, C.M., Bao, H., Hendriks, I.A., Carraro, M., García-Nieto, A., Liu, Y., Reverón-Gómez, N., Spanos, C., Chen, L., Rappsilber, J., et al. (2021) DNAJC9 integrates heat shock molecular chaperones into the histone chaperone network. Mol. Cell, 81, 2533–2548.e9.

33. Adam, S., Dabin, J., Bai, S.-K. and Polo, S.E. (2015) Imaging local deposition of newly synthesized histones in UVC-damaged chromatin. *Methods Mol. Biol. (Clifton*, NJ), 1288, 337–47.

34. Ray-Gallet, D., Woolfe, A., Vassias, I., Pellentz, C., Lacoste, N., Puri, A., Schultz, D.C., Pchelintsev, N.A., Adams, P.D., Jansen, L.E.T., et al. (2011) Dynamics of Histone H3 Deposition In Vivo Reveal a Nucleosome Gap-Filling Mechanism for H3.3 to Maintain Chromatin Integrity. Mol. Cell, 44, 928–941.

35. Reid, D.A., Reed, P.J., Schlachetzki, J.C.M., Nitulescu, I.I., Chou, G., Tsui, E.C., Jones, J.R., Chandran, S., Lu, A.T., McClain, C.A., et al. (2021) Incorporation of a nucleoside analog maps genome repair sites in postmitotic human neurons. Science, 372, 91–94.

36. Carraro, M., Hendriks, I.A., Hammond, C.M., Solis-Mezarino, V., Völker-Albert, M., Elsborg, J.D., Weisser, M.B., Spanos, C., Montoya, G., Rappsilber, J., et al. (2023) DAXX adds a de novo H3.3K9me3 deposition pathway to the histone chaperone network. Mol. Cell, 83, 1075–1092.e9.

37. Smith, C.L., Matheson, T.D., Trombly, D.J., Sun, X., Campeau, E., Han, X., Yates, J.R. and Kaufman, P.D. (2014) A separable domain of the p150 subunit of human Chromatin Assembly Factor-1 promotes protein and chromosome associations with nucleoli. Mol. Biol. Cell, 25, mbc.E14-05-1029.

38. Balachandra, V., Shrestha, R.L., Hammond, C.M., Lin, S., Hendriks, I.A., Sethi, S.C., Chen, L., Sevilla, S., Caplen, N.J., Chari, R., et al. (2024) DNAJC9 prevents CENP-A mislocalization and chromosomal instability by maintaining the fidelity of histone supply chains. EMBO J., 43, 2166–2197.

39. Wessel, S.R., Mohni, K.N., Luzwick, J.W., Dungrawala, H. and Cortez, D. (2019) Functional Analysis of the Replication Fork Proteome Identifies BET Proteins as PCNA Regulators. Cell Rep., 28, 3497–3509.e4.

40. Marteijn, J.A., Lans, H., Vermeulen, W. and Hoeijmakers, J.H.J. (2014) Understanding nucleotide excision repair and its roles in cancer and ageing. Nat. Rev. Mol. Cell Biol., 15, 465–481.

41. Sugasawa, K., Ng, J.M.Y., Masutani, C., Maekawa, T., Uchida, A., Spek, P.J. van der Eker, A.P.M., Rademakers, S., Visser, C., Aboussekhra, A., et al. (1997) Two Human Homologs of Rad23 Are Functionally Interchangeable in Complex Formation and Stimulation of XPC Repair Activity. Mol. Cell. Biol., 17, 6924–6931.

42. Horianopoulos, L.C., Lee, C.W.J., Schmitt, K., Valerius, O., Hu, G., Caza, M., Braus, G.H. and Kronstad, J.W. (2021) A J Domain Protein Functions as a Histone Chaperone to Maintain Genome Integrity and the Response to DNA Damage in a Human Fungal Pathogen. mBio, 12, e03273–21.

43. Lemaître, C., Grabarz, A., Tsouroula, K., Andronov, L., Furst, A., Pankotai, T., Heyer, V., Rogier, M., Attwood, K.M., Kessler, P., et al. (2014) Nuclear position dictates DNA repair pathway choice. Genes Dev., 28, 2450–2463.

44. Lemaître, C., Fischer, B., Kalousi, A., Hoffbeck, A.-S., Guirouilh-Barbat, J., Shahar, O.D., Genet, D., Goldberg, M., Betrand, P., Lopez, B., et al. (2012) The nucleoporin 153, a novel factor in double-strand break repair and DNA damage response. Oncogene, 31, 4803– 4809.

45. Ryu, T., Spatola, B., Delabaere, L., Bowlin, K., Hopp, H., Kunitake, R., Karpen, G.H. and Chiolo, I. (2015) Heterochromatic breaks move to the nuclear periphery to continue recombinational repair. Nat. Cell Biol., 17, 1401–1411.

46. Schep, R., Brinkman, E.K., Leemans, C., Vergara, X., Weide, R.H. van der, Morris, B., Schaik, T. van Manzo, S.G., Peric-Hupkes, D., Berg, J. van den et al. (2021) Impact of chromatin context on Cas9-induced DNA double-strand break repair pathway balance. Mol. Cell, 81, 2216–2230.e10.

47. Dunleavy, E.M., Almouzni, G. and Karpen, G.H. (2011) H3.3 is deposited at centromeres in S phase as a placeholder for newly assembled CENP-A in G1 phase. Nucleus, 2, 146–157.

48. Katsumi, S., Kobayashi, N., Imoto, K., Nakagawa, A., Yamashina, Y., Muramatsu, T., Shirai, T., Miyagawa, S., Sugiura, S., Hanaoka, F., et al. (2001) In Situ Visualization of Ultraviolet-Light-Induced DNA Damage Repair in Locally Irradiated Human Fibroblasts. J. Investig. Dermatol., 117, 1156–1161.

49. Moné, M.J., Volker, M., Nikaido, O., Mullenders, L.H.F., Zeeland, A.A. van, Verschure, P.J., Manders, E.M.M. and Driel, R. van (2001) Local UV-induced DNA damage in cell nuclei results in local transcription inhibition. EMBO Rep., 2, 1013–1017.

50. Dinant, C., Jager, M. de, Essers, J., Cappellen, W.A. van, Kanaar, R., Houtsmuller, A.B. and Vermeulen, W. (2007) Activation of multiple DNA repair pathways by sub-nuclear damage induction methods. J. Cell Sci., 120, 2731–2740.

51. Casabona, M.G., Vandenbrouck, Y., Attree, I. and Couté, Y. (2013) Proteomic characterization of Pseudomonas aeruginosa PAO1 inner membrane. PROTEOMICS, 13, 2419–2423.

52. Bouyssié, D., Hesse, A.-M., Mouton-Barbosa, E., Rompais, M., Macron, C., Carapito, C., Peredo, A.G. de, Couté, Y., Dupierris, V., Burel, A., et al. (2020) Proline: an efficient and user-friendly software suite for large-scale proteomics. Bioinformatics, 36, 3148–3155.

53. Couté, Y., Bruley, C. and Burger, T. (2020) Beyond Target–Decoy Competition: Stable Validation of Peptide and Protein Identifications in Mass Spectrometry-Based Discovery Proteomics. Anal. Chem., 92, 14898–14906.

54. Perez-Riverol, Y., Csordas, A., Bai, J., Bernal-Llinares, M., Hewapathirana, S., Kundu, D.J., Inuganti, A., Griss, J., Mayer, G., Eisenacher, M., et al. (2018) The PRIDE database and related tools and resources in 2019: improving support for quantification data. Nucleic Acids Res., 47, gky1106-.

55. Wieczorek, S., Combes, F., Lazar, C., Gianetto, Q.G., Gatto, L., Dorffer, A., Hesse, A.-M., Couté, Y., Ferro, M., Bruley, C., et al. (2017) DAPAR & ProStaR: software to perform statistical analyses in quantitative discovery proteomics. Bioinformatics, 33, 135–136.

56. Medvedeva, Y.A., Lennartsson, A., Ehsani, R., Kulakovskiy, I.V., Vorontsov, I.E., Panahandeh, P., Khimulya, G., Kasukawa, T., Consortium, T.F. and Drabløs, F. (2015) EpiFactors: a comprehensive database of human epigenetic factors and complexes. Database, 2015, bav067.

57. Marakulina, D., Vorontsov, I.E., Kulakovskiy, I.V., Lennartsson, A., Drabløs, F. and Medvedeva, Y.A. (2022) EpiFactors 2022: expansion and enhancement of a curated database of human epigenetic factors and complexes. Nucleic Acids Res., 51, D564–D570.

58. Bodor, D.L., Rodríguez, M.G., Moreno, N. and Jansen, L.E.T. (2012) Analysis of Protein Turnover by Quantitative SNAP-Based Pulse-Chase Imaging. Curr. Protoc. Cell Biol., 55, 8.8.1–8.8.34.

