## Supplementary figures for "Proteomic profiling of UV damage repair patches uncovers histone chaperones with central functions in chromatin repair"

### Supplementary figure 1

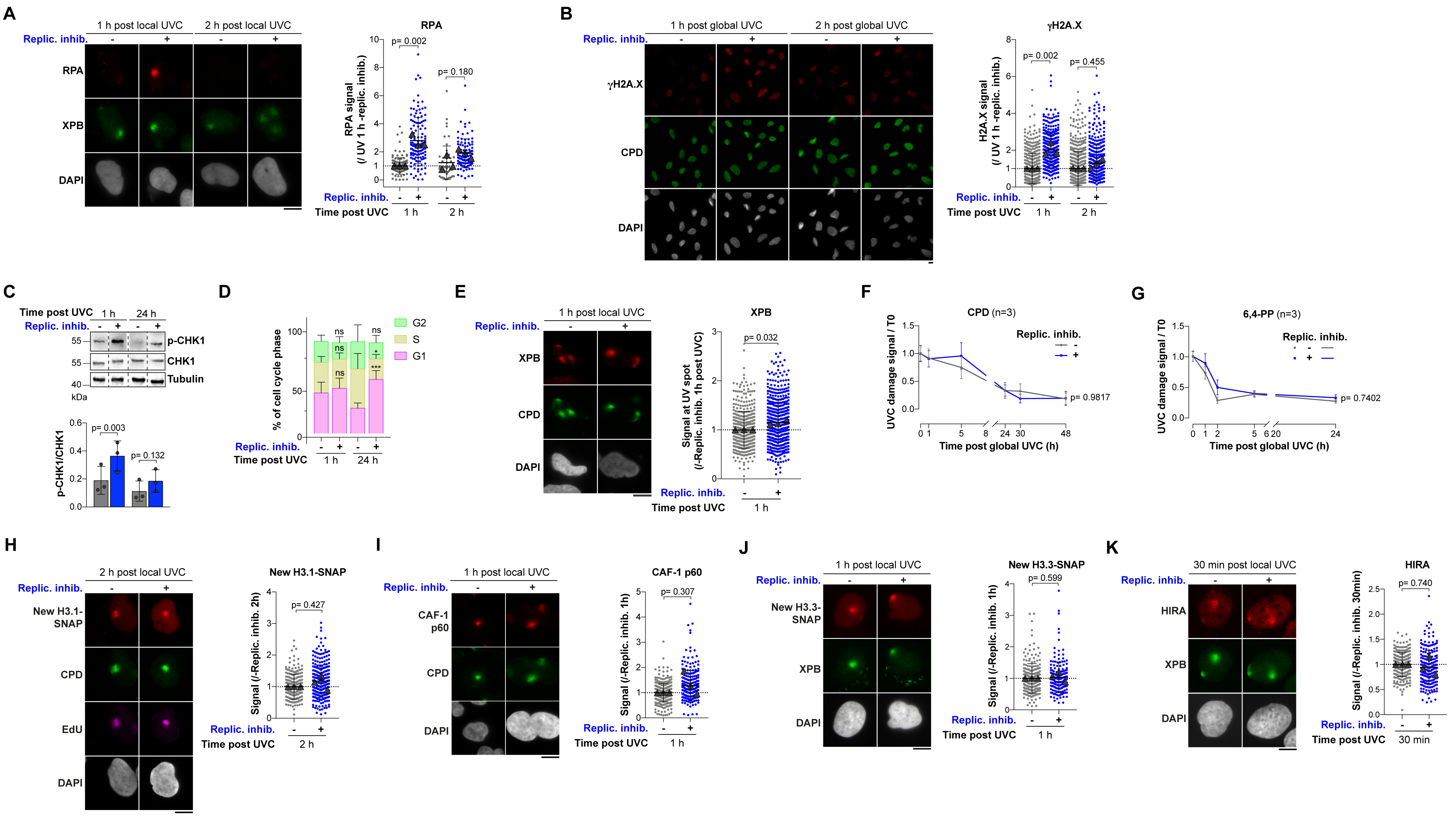

### Supplementary figure 2

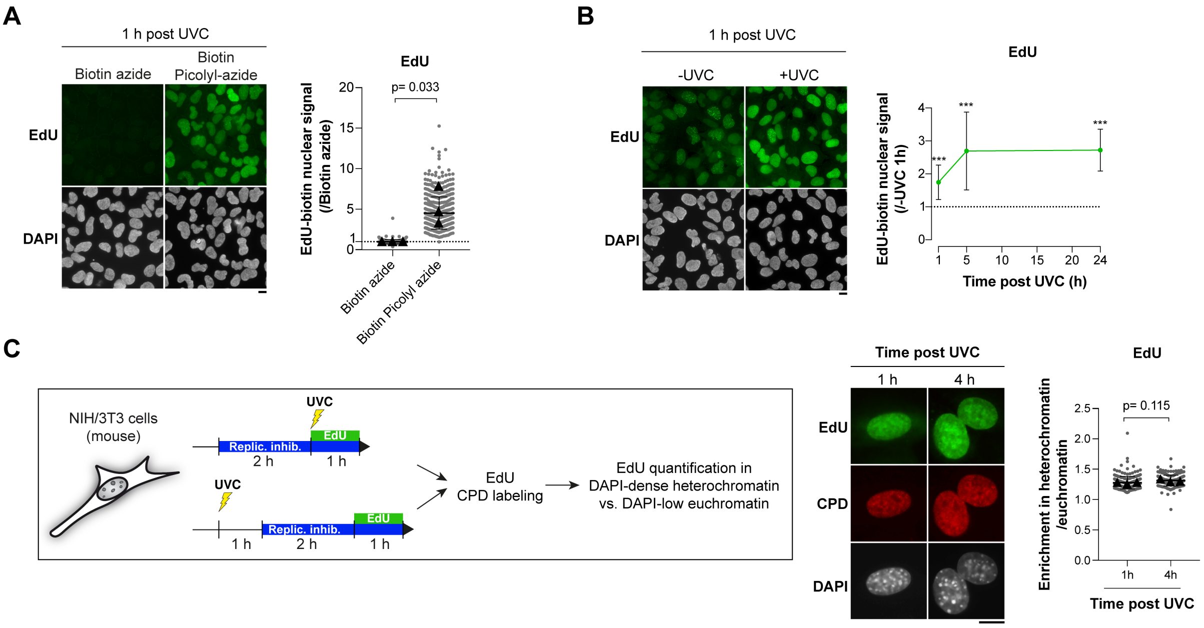

#### Supplementary figure 3

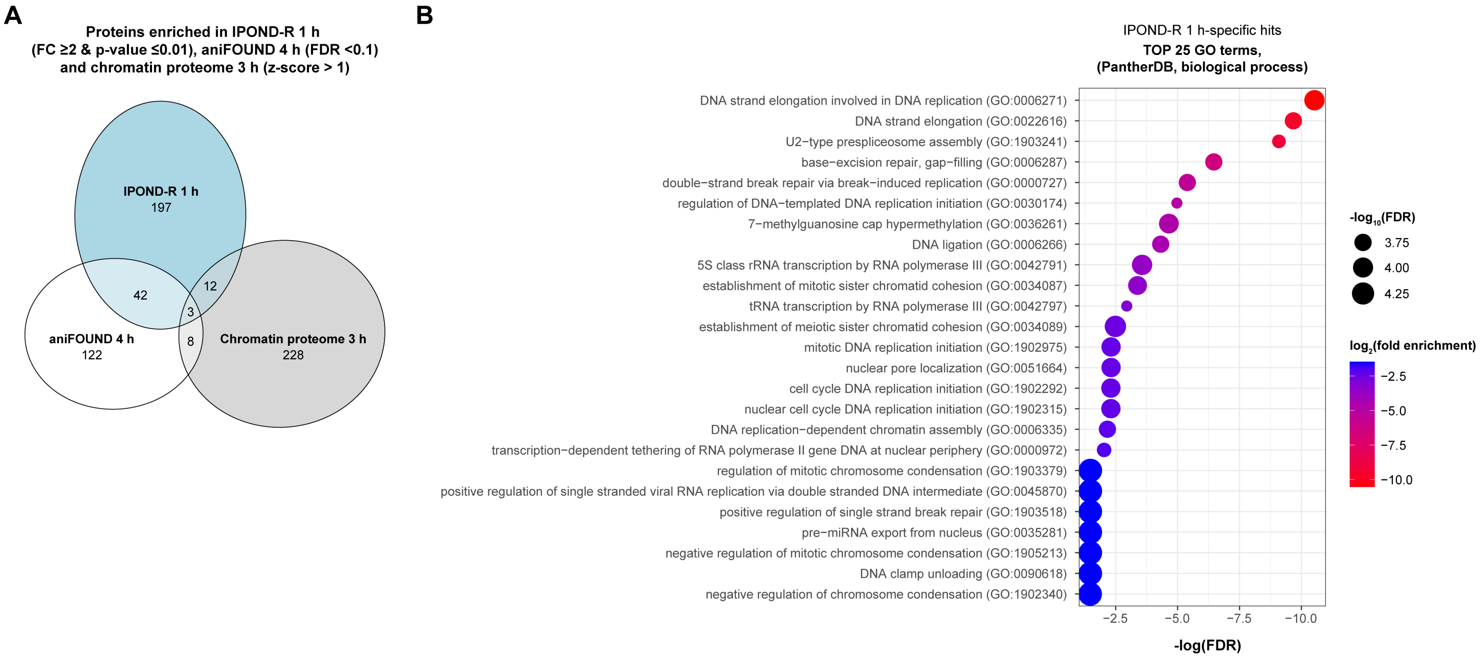

#### Supplementary figure 4

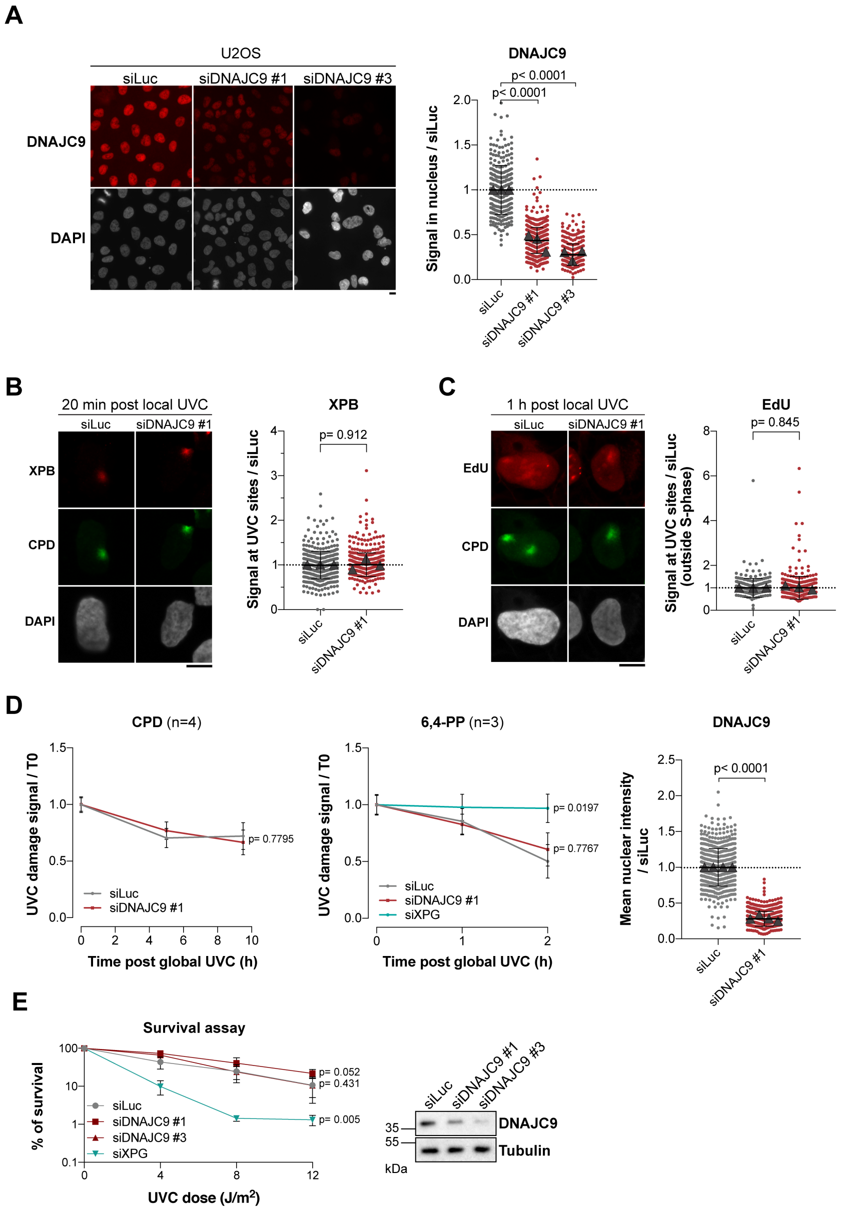

#### Supplementary figure 5

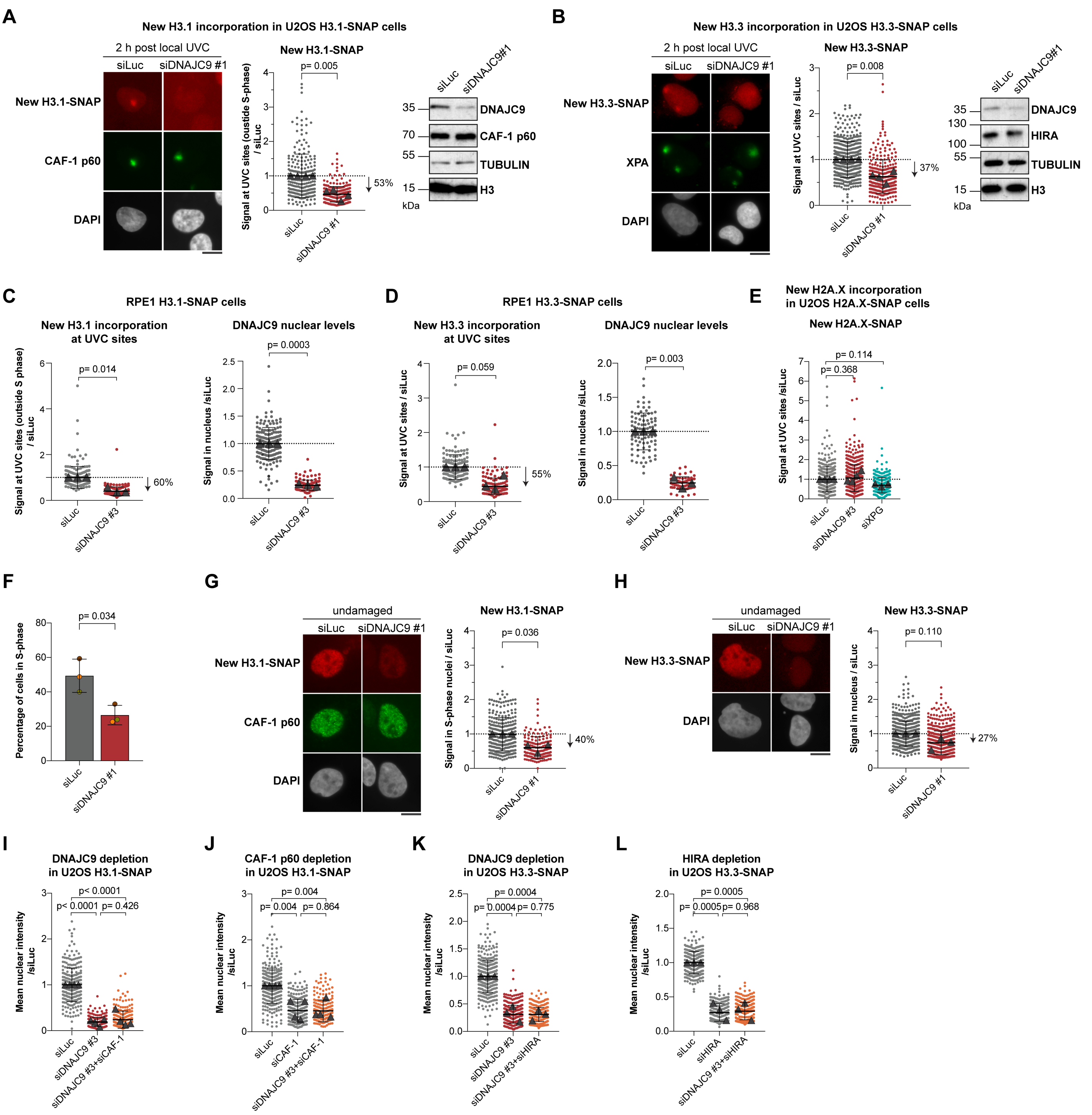

### Supplementary figure 6

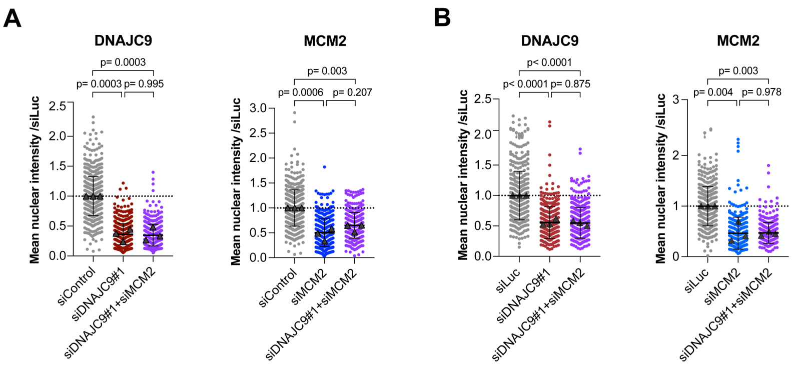
